# Neuronal ribosomes dynamically exchange ribosomal proteins in a context-dependent manner

**DOI:** 10.1101/2021.03.25.437026

**Authors:** Claudia M. Fusco, Kristina Desch, Aline R. Dörrbaum, Mantian Wang, Anja Staab, Ivy C.W. Chan, Eleanor Vail, Veronica Villeri, Julian D. Langer, Erin M. Schuman

## Abstract

Owing to their morphological complexity and dense network connections, neurons modify their proteomes locally, using mRNAs and ribosomes present in the neuropil (tissue enriched for dendrites and axons). Although ribosome biogenesis largely takes place in the nucleus and perinuclear region, neuronal ribosomal protein (RP) mRNAs have been frequently detected remotely, in dendrites and axons. Here, using imaging and ribosome profiling, we directly detected the RP mRNAs and their translation in the neuropil. Combining brief metabolic labeling with mass spectrometry, we found that a group of RPs quickly associated with translating ribosomes in the cytoplasm and that this incorporation is independent of canonical ribosome biogenesis. Moreover, the incorporation probability of some RPs was regulated by location (neurites vs. cell bodies) and changes in the cellular environment (in response to oxidative stress). Our results suggest new mechanisms for the local activation, repair and/or specialization of the translational machinery within neuronal processes, potentially allowing remote neuronal synapses a rapid solution to the relatively slow and energy-demanding requirement of nuclear ribosome biogenesis.

## INTRODUCTION

Neurons use the translation of distally localized mRNAs for synapse formation, axon growth and synaptic plasticity (Holt et al., 2019). Although ribosomes have been detected in dendrites (Bodian, 1965) and axons (Hafner et al., 2019; Shigeoka et al., 2019; Tennyson, 1970), little is known about ribosome biogenesis and homeostasis in neurons. Thus far, studies from yeast to human have revealed a striking conservation of the basic ribosome structure (Anger et al., 2013). Eukaryotic ribosomes are composed of a small and a large subunit comprising ∼79 proteins (ribosomal proteins, RPs) and 4 rRNA species. In eukaryotes, ribosomal components (including most RPs and rRNA) are initially co-assembled in the nucleolus. The nearly mature ribosome is then exported to the cytoplasm where a few RPs associate to complete the maturation process (la Cruz et al., 2015). RPs are thought to exhibit a stable, “life-long” incorporation with their associated subunits, and, at the end of their life-cycle, to undergo concerted degradation (An and Harper, 2019).

Recent data, however, have suggested that ribosomes may be less static than the above picture suggests (Emmott et al., 2018; Genuth and Barna, 2018). Proteomic data, for example, have reported ribosomes containing individual ribosomal proteins at different stoichiometries with unique translational properties (Shi et al., 2017; Slavov et al., 2015). As ribosome biogenesis is believed to require the step-wise incorporation of all ribosomal proteins, it remains unclear how heterogeneous ribosomes are formed. Additionally, many transcriptomics studies have detected RP mRNAs remote from the site of ribosome biogenesis, including in distal neuronal processes, potentially challenging our common understanding of assembled ribosome as a static structure (Andreassi et al., 2010; Biever et al., 2020; Briese et al., 2016; Cajigas et al., 2012; Gioio et al., 2004; Hafner et al., 2019; Mardakheh et al., 2015; Mazaré et al., 2020; Middleton et al., 2019; Misra et al., 2016; Moccia et al., 2003a; Moor et al., 2017; Perez et al., 2021; Poulopoulos et al., 2019; Saal et al., 2014; Shigeoka et al., 2016; Taylor et al., 2009; Tushev et al., 2018; Zivraj et al., 2010). Indeed, the cytosolic incorporation of some RPs in developing Xenopus retinal ganglion cell axons was recently observed (Shigeoka et al., 2019). Altogether, the above data suggest that the protein composition of ribosomes might not be fixed after biogenesis, but rather be subject to dynamic association or exchange of nascent RPs with mature ribosomes.

To address this possibility in neurons, we first used high-resolution fluorescence *in situ* hybridization (FISH) to directly detect a large population of RP mRNAs in rodent neuronal cell bodies and dendrites. Using ribosome footprinting and metabolic-labeling approaches we observed the active translation of RP mRNAs in the neuropil. The dendritic synthesis of RPs, remote from the peri-nuclear region, prompted us to investigate the dynamics of RP association with mature neuronal ribosomes. We used very brief metabolic labeling (pSILAC) combined with parallel-reaction monitoring mass spectrometry to evaluate selectively the abundance of individual “new” and “old” RP peptides within ribosomes. We identified a population of 12 nascent RPs (“exchangers”) that rapidly incorporate into mature pre-existing ribosomes. Using compartmentalized chambers, we observed the biogenesis-independent incorporation of RPs in both somata and isolated neuronal processes. Moreover, we found that the incorporation probability of some RPs was regulated by the subcellular compartment (neurites vs. cell bodies) and by changes in the physiological state (e.g. during oxidative stress). Taken together, these data suggest that neurons can dynamically regulate RPs incorporation into ribosomes in space and time.

## RESULTS

### RP mRNA localization and translation in dendrites

Advances in transcriptome-wide profiling methods have led to the elucidation of thousands of mRNAs localized to neuronal processes. In addition to many neuronal/synaptic transcripts, the RP mRNAs have been surprisingly detected in many preparations enriched for axons and dendrites (Figure 1A and Table S1; see also (Andreassi et al., 2010; Biever et al., 2020; Briese et al., 2016; Cajigas et al., 2012; Gioio et al., 2004; Gumy et al., 2011; Hafner et al., 2019; Middleton et al., 2019; Moccia et al., 2003b; Perez et al., 2021; Poulopoulos et al., 2019; Saal et al., 2014; Shigeoka et al., 2016; Taylor et al., 2009; Tushev et al., 2018; Zivraj et al., 2010). To evaluate whether the mRNAs for the ribosome are specifically enriched in dendrites and axons, we compared the dendritic enrichment of RP mRNAs to mRNAs that code for proteins in other ubiquitous macromolecular complexes. We used total RNA-seq data (Biever et al., 2020), comparing somata-enriched or neuropil fractions of rat hippocampal slices (Figure 1B) and quantified the neuropil enrichment of mRNAs coding proteins of the ribosome (RPs), proteasome, nuclear pore complex and RNA polymerase I-III (Figure 1C). We found that only the RP mRNAs exhibited a consistent enrichment in the neuropil, while the mRNAs of all other complexes were mostly enriched in somata. This suggests that the neuropil localization of RP mRNAs is not owing to “background” detection of abundant mRNAs or the presence of contaminants in the sequenced material.

**Figure 1.**
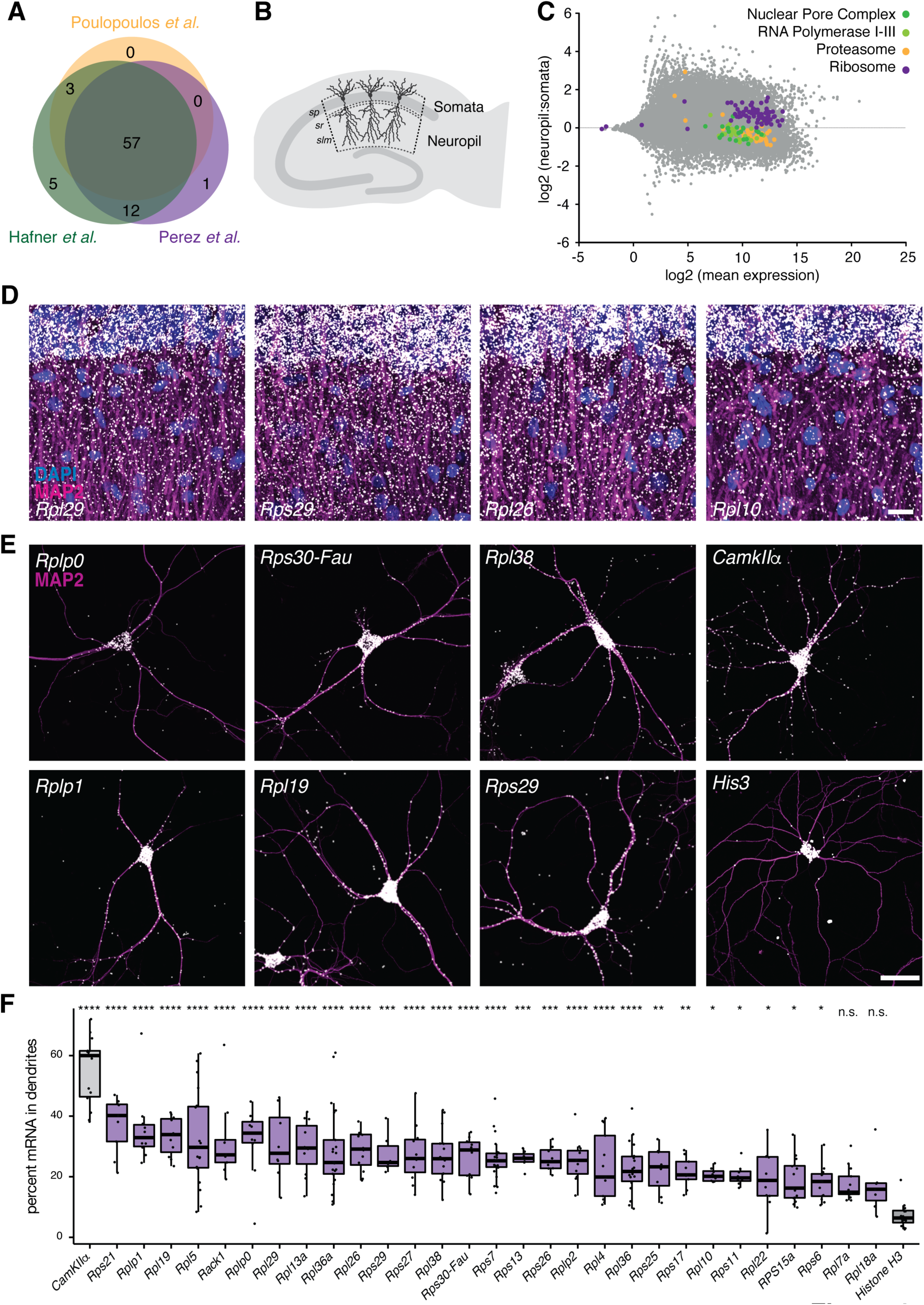
RP mRNAs are present in hippocampal dendrites. (A) Overlap of RP mRNAs detected in neuronal processes from three different studies: dendritic arbors ((Perez et al., 2021), in purple), purified synaptosomes ((Hafner et al., 2019), in green), developing axons ((Poulopoulos et al., 2019), in yellow) of rodent neurons. See also Table S1 for additional studies and data. (B) Schematic representation of a hippocampal slice. Three layers in area CA1 are shown: somata (*stratum pyramidale*, rich in cell bodies) and neuropil (*strata radiatum et lacunosum-moleculare*, rich in axons and dendrites (Mishchenko et al., 2010). (C) MA plot depicting the expression of mRNAs from the somata and neuropil of hippocampal slices (Biever et al., 2020). Each mRNA is represented by a single data point. The y-axis depicts the relative expression in the neuropil or somata and the x-axis depicts the mean expression of each mRNA. Unlike mRNAs for other macro-molecular complexes (e.g. the proteasome, nuclear pore complex and RNA polymerase I-III), mRNAs coding for ribosomal proteins are consistently enriched in the neuropil. (D) FISH detection of indicated RP mRNAs in dendrites (magenta MAP2, white FISH, blue DAPI) in hippocampal slices. Images are oriented with the somata layer at the top and the dendrites extending below. Scale bar = 50 µm. See Figure S1a-b for additional RPs and analysis. (E) FISH detection of indicated RP mRNAs as well as CamkIIα and His3 mRNA (magenta MAP2, white FISH) in cultured hippocampal neurons Scale bar = 50 µm. See Figure S2 for additional RPs. (F) Analysis of FISH data shown in E and Figure S2. Percentage of indicated mRNA signal in dendrites over the total detected in single neurons. Transcripts are ranked according to average value. The distribution of each mRNA was compared to the largely somatically-localized transcript *His3*. ANOVA (p<0.0001) followed by a Dunnett’s multiple comparison test, ** p≤0.01, *** p≤0.001, **** p≤0.0001.

To assess directly whether the RP mRNAs are localized in dendrites, we performed single molecule fluorescence *in situ* hybridization (smFISH) for 29 different endogenous RP transcripts in both rat hippocampal slices (Figure 1D and Figure S1A) and cultured rat hippocampal neurons (Figure 1E and Figure S2). In hippocampal slices, for each RP transcript evaluated, we detected signal in the somata (*s. pyramidale*) and in the neuropil (*s. radiatum*) at levels similar to those measured by RNAseq (Figure S1B-C). Likewise, in cultured hippocampal neurons, we detected abundant RP mRNAs both in the cell body as well as in the dendrites. To normalize for potential differences in expression level, we quantified the fraction of the total mRNA signal detected in the dendrites of individual neurons. Amongst the 29 RP mRNAs we evaluated, 15% (e.g. Rpl7a) to 40% (e.g. Rps21) of the total mRNA was localized in dendrites (Figure 1F). For comparison we analyzed the dendritic abundance of a well-studied and abundant dendritic mRNA Ca^2+^-calmodulin-dependent protein kinase, CamKIIα (Burgin et al., 1990; Cajigas et al., 2012; Miller et al., 2002), and a somatic-enriched mRNA encoding the nuclear protein histone H3-3B (Cajigas et al., 2012). As expected, a high fraction (60%) of CamKIIα mRNAs and a very low fraction (6%) of Histone 3 mRNAs were detected in dendrites. Each of the 29 tested RP mRNAs exhibited a distribution in the dendrites greater than that observed for the nuclear protein encoding mRNA Histone H3 (Fig.1F). Taken together, these data demonstrate that RP mRNAs are localized to the neuropil and dendrites of hippocampal slices and cultured neurons.

We next asked whether the dendritically localized RP mRNAs are locally translated into protein. We investigated via ribosome profiling whether RP mRNAs are associated with translating ribosomes in the cell bodies and/or neuropil of the hippocampus. Using our dataset (Biever et al., 2020), we detected ribosome footprints across the entire coding sequence of each RP transcript measured in the neuropil, a region enriched for axons and dendrites (Figure 2A and Figure S3). Analysis of the footprint abundance revealed that all RP mRNAs were either equally translated within the two compartments (somata or neuropil) or exhibited significantly enhanced translation in the neuropil (Figure S4A). We note that the translation of RPs has also been recently reported in mouse retinal ganglion cell axons (Cagnetta et al., 2018; Shigeoka et al., 2016). Furthermore, using puromycin proximity ligation assay (Puro-PLA) to visualize newly synthesized proteins-of-interest (tom Dieck et al., 2015), we observed nascent signal within the dendrites for all 17 RPs examined with just 5 min of metabolic labeling (Figure 2B and Figure S5). For example, almost half of the total RPL19 signal was observed in the dendrites (Figure S4C). As expected, the addition of a protein synthesis inhibitor significantly inhibited the dendritic nascent protein signal of all RPs (Figures 2B-C, S4B and S5). Moreover, the nascent dendritic protein signal did not increase when a 5 min “chase” followed the metabolic label (Figure S4D-E), suggesting that, over short time scales, there was no significant contribution of somatically synthesized RPs to the measured dendritic nascent protein signal. Taken together, these data indicate that RP mRNAs are locally translated in the neuropil and dendrites of hippocampal neurons.

**Figure 2.**
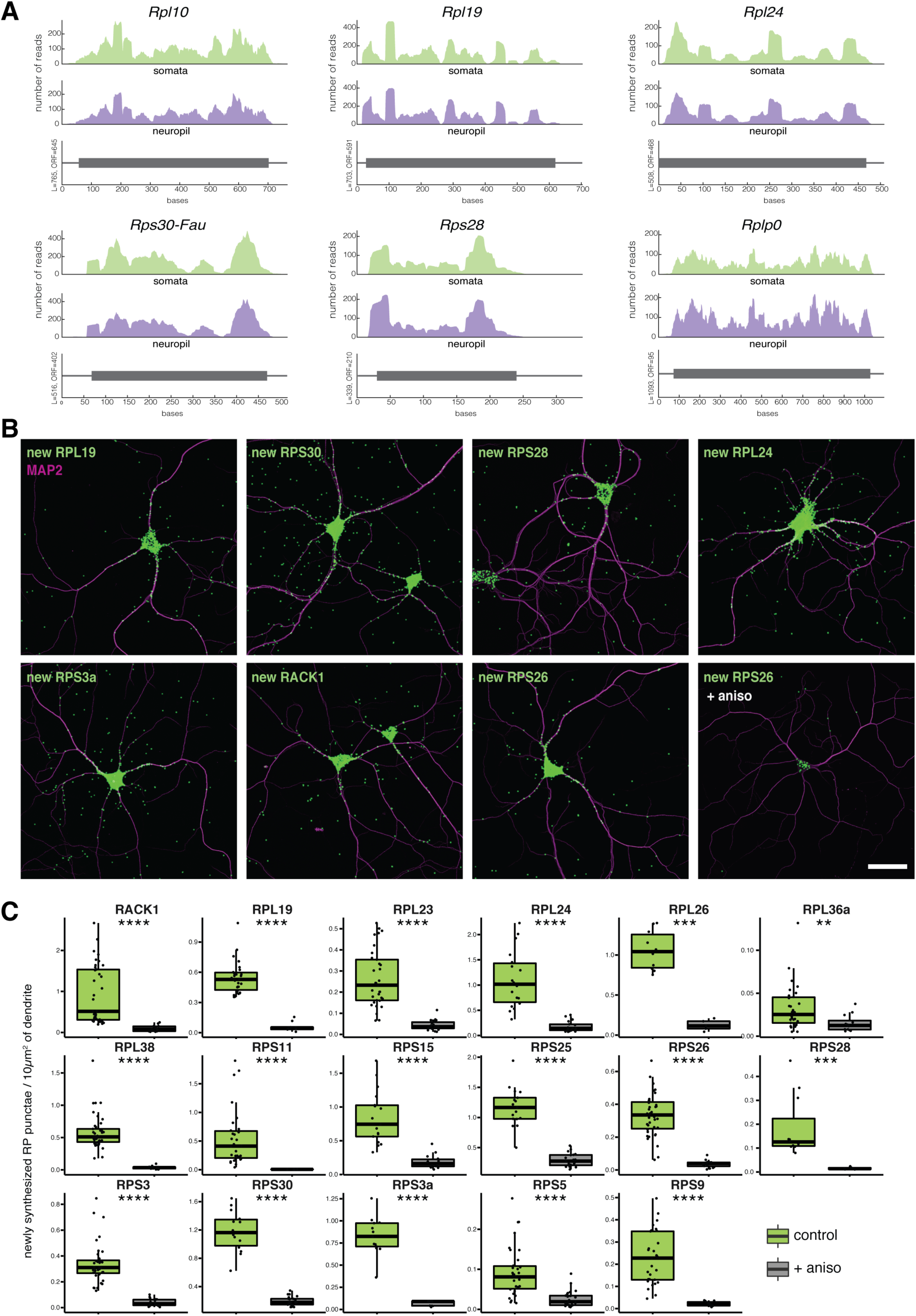
RP mRNAs are translated in hippocampal dendrites. (A) Ribosome footprint coverage of RP mRNAs from the somata and neuropil of hippocampal slices (Biever et al., 2020). Shown are the number of reads throughout the open reading frame (grey box) from the somata-enriched fraction (green) or the neuropil-enriched fraction (purple). Ribosome footprint coverage for additional RPs is shown in Figure S3. (B) Detection of nascent RPs (green) in dendrites (magenta for MAP2 immunostaining) of cultured hippocampal neurons using Puro-PLA (tom Dieck et al., 2015). Nascent proteins were labeled with puromycin (5 min), in the absence or presence (as indicated) of the protein synthesis inhibitor anisomycin (see methods). Scale bar = 50 µm. Data for additional nascent RP detection are shown in Figures S4B and 5. (C) Analysis of nascent RP detection shown in Figures 2B, S4B and 5. Analyzed are the newly synthesized RP punctae per 10µm^2^ of dendrite, in control (green) or in the presence of protein synthesis inhibitor (grey). Each dot is the quantification from the whole dendritic arbor of one single neuron. Wilcox test, ** p≤0.01, *** p≤0.001, **** p≤0.0001.

### Dynamic association of nascent RPs with mature ribosomes

The dendritic synthesis of RPs, remote from the peri-nuclear region, prompted us to investigate the dynamics of RP association with mature neuronal ribosomes. We followed the incorporation kinetics of individual RPs into assembled ribosomes, asking whether all individual RPs are incorporated at an equal rate or whether there is a sub-population that is incorporated with different kinetics. To do so, we metabolically labeled newly synthesized proteins by incubating neurons for 1 or 2 hours with heavy amino acids (pSILAC, (Bogenhagen et al., 2018; Ross et al., 2021; Schwanhäusser et al., 2009)). We then purified translating ribosomes using a sucrose cushion (modified from (McGlincy and Ingolia, 2017)) and used mass spectrometry to quantify the fraction of new RPs present in assembled ribosomes (Figure 3A). A sample without heavy amino acids served as negative control. We verified the translating status of the ribosomes by confirming the sensitivity of our preparation to conditions that disassemble monosomes and polysomes into free small and large subunits (in the absence of magnesium or in the presence of the chelating agent EDTA, Figure S6A-B). Note that as the average half-life of a brain RP is ∼8 days (Dörrbaum et al., 2018; Fornasiero et al., 2018; Stoykova et al., 1983), only ∼0.4% of each protein is expected to be synthesized after 1 hour of labeling (Figure S6C). Using a targeted mass spectrometry method (Parallel Reaction Monitoring (Peterson et al., 2012)) to maximize our sensitivity, we reliably quantified 70 new RPs after 1 or 2 hrs of labeling (Figure 3B and S6D). For all RPs, we observed an increase in nascent RP incorporation into assembled ribosomes with increased labeling time (1 vs 2 hrs) and the labeling level of individual RPs was significantly correlated between the 2 timepoints (r^2^ = 0.84, p < 0.0001; Figure 3C). Importantly, the omission of the heavy amino acids led to a complete loss of the heavy peptide peak at the expected position (Figure S6D). In addition, there was no significant difference in the ribosome association between nascent RPs that comprise large and small ribosome subunits (Figure S6E-G). Interestingly, however, individual nascent RPs exhibited distinct kinetics of accumulation in mature ribosomes, with the labeling fraction varying by >3.5-fold (Figure 3B-C-E). To identify RPs that share similar kinetics we performed an unsupervised hierarchical clustering and detected 6 RP groups (Figure 3B). Three groups (comprising a total of 12 proteins) exhibited a higher association level with assembled ribosomes (“rapidly incorporating”, clusters A-B-C in Figure 3B-F) than the other 3 groups of RPs (comprising 58 proteins) (clusters D-E-F in Figure 3B-F). This higher association rate of the rapidly incorporating group was detected following both 1 and 2 hrs of labeling (Figure 3B-D). Within this group, we noted the presence of several RPs known to associate late during ribosome biogenesis, like RACK1 and RPL10 (la Cruz et al., 2015; Larburu et al., 2016) validating the sensitivity of our measurements to established temporal dynamics of ribosome assembly (Figure S6H-I).

**Figure 3.**
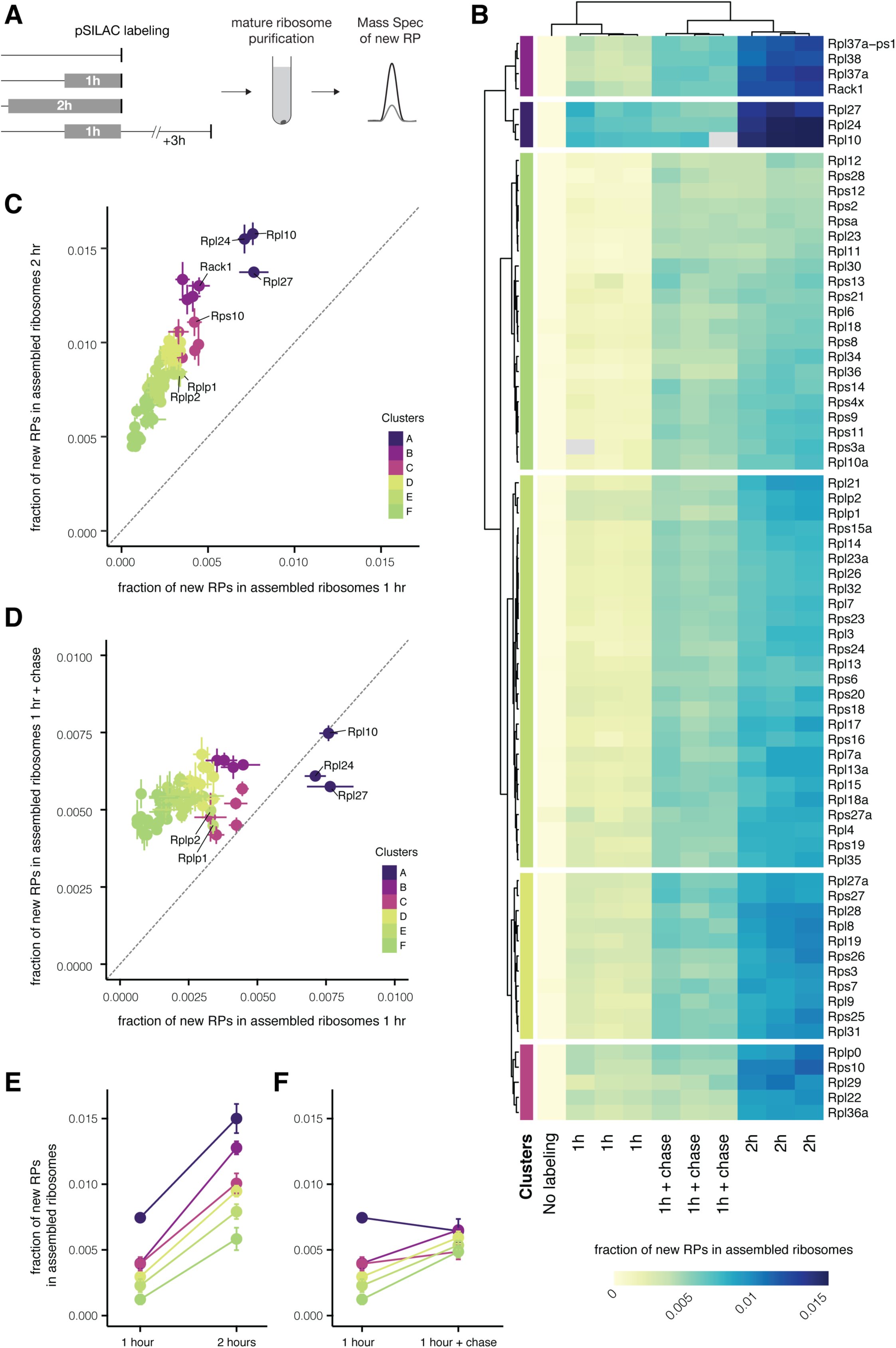
Some RPs rapidly incorporate into neuronal ribosome. (A) Schematic of the experimental design for measuring new RPs in assembled ribosomes from cultured neurons. Normal medium was replaced by medium containing heavy amino acids (pSILAC) as indicated (grey boxes). Mature ribosomes were purified by sucrose cushion. New ribosomal proteins were quantified by mass spectrometry, measuring the heavy and light peak of each peptide. (B) Heatmap showing for each RP the fraction of new proteins (H/(H+L)) incorporated into assembled ribosomes. Pseudocells (median of peptides obtained per individual protein) are ordered according to unsupervised clustering, both for columns (biological replicate of each condition) and rows (individual ribosomal protein). Experimental conditions of the labeling are indicated at the bottom. (C-D) Scatterplots showing the fraction of new RPs (H/(H+L)) in assembled ribosomes after the different labeling conditions, as indicated by x- and y- axes. Points represent average +/- standard deviation of three biological replicates. Proteins are colored according to clusters identified in Figure 3B. Some RPs of interest are indicated by name. Dashed line represents x=y. (E-F) Average +/- standard deviation of the fraction of new proteins in assembled ribosomes of RPs of the same cluster, as identified in Figure 3B. The different labeling conditions are indicated on the x-axis.

To measure the potential time-lag between an individual RP’s synthesis and its association with mature ribosomes we performed a pulse-chase experiment. We labeled nascent RPs for 1 hr and imposed a 3 hrs (label-free) chase before the purification of assembled ribosomes (Figure 3A). Consistent with a time delay between synthesis and incorporation, the addition of the chase period led to a selective increase in mature ribosome association for the nascent RPs that showed slower incorporation kinetics (clusters D-E-F) in the previous 1 and 2 hrs labeling experiments (Figure 3B-D-F). Interestingly, the remaining clusters (A-C) exhibited lower levels of incorporation after the chase. RPL27, RPL10 and RPL24 (cluster A), which showed the highest incorporation after 1 hr labeling (no-chase, Figure 3D), actually decreased their association with mature ribosomes when the chase was imposed (Figure 3F). This indicates that either they rapidly and persistently associate with mature ribosomes or that they are replaced (exchanged) by nascent (but un-labeled) proteins synthesized during the chase. Supporting the idea of RP exchange, we noted the presence of RPLP1 and 2 among the proteins with the lowest fold change after the chase; these proteins are the only two known RPs to transiently associate and dissociate from mature ribosomes (Tsurugi and Ogata, 1985; Zinker, 2014). Taken together, these data reveal that the binding kinetics of RPs to neuronal ribosomes are not homogeneous, and that a subset of RPs can rapidly and dynamically incorporate into neuronal ribosomes. Importantly, we note that the 12/12 mRNAs of the rapidly incorporating RPs were detected in the neuropil RNA-seq dataset (Figure 1C), and 11/12 were detected in the ribosome footprints dataset (Figure 2A and S3). Additionally, we visualized the dendritic localization of both the mRNA (Figure 1E and Figure S2) and the nascent protein (Figure 2B and Figure S5), for all tested members of this group.

### Biogenesis-independent incorporation of nascent RPs

To test whether the above observed dynamic incorporation might represent the exchange of some nascent RPs on mature ribosomes, we determined the sensitivity of the rapid RP association to inhibition of ribosome biogenesis. We used leptomycin B (LMB) to block nuclear export (Thomas and Kutay, 2003); resulting in an inhibition of ribosome biogenesis. The inhibition of nuclear export (ribosome biogenesis) by LMB was confirmed by the following: i) the inhibition of CMR1-mediated nuclear transport (Figure S7A-B), resulting in a sequestration of RPs (Figure S7C-D) and rRNA (Figure S7E-F) in the nucleus, ii) the depletion of nascent rRNA in assembled ribosomes (Figure 4B).

**Figure 4.**
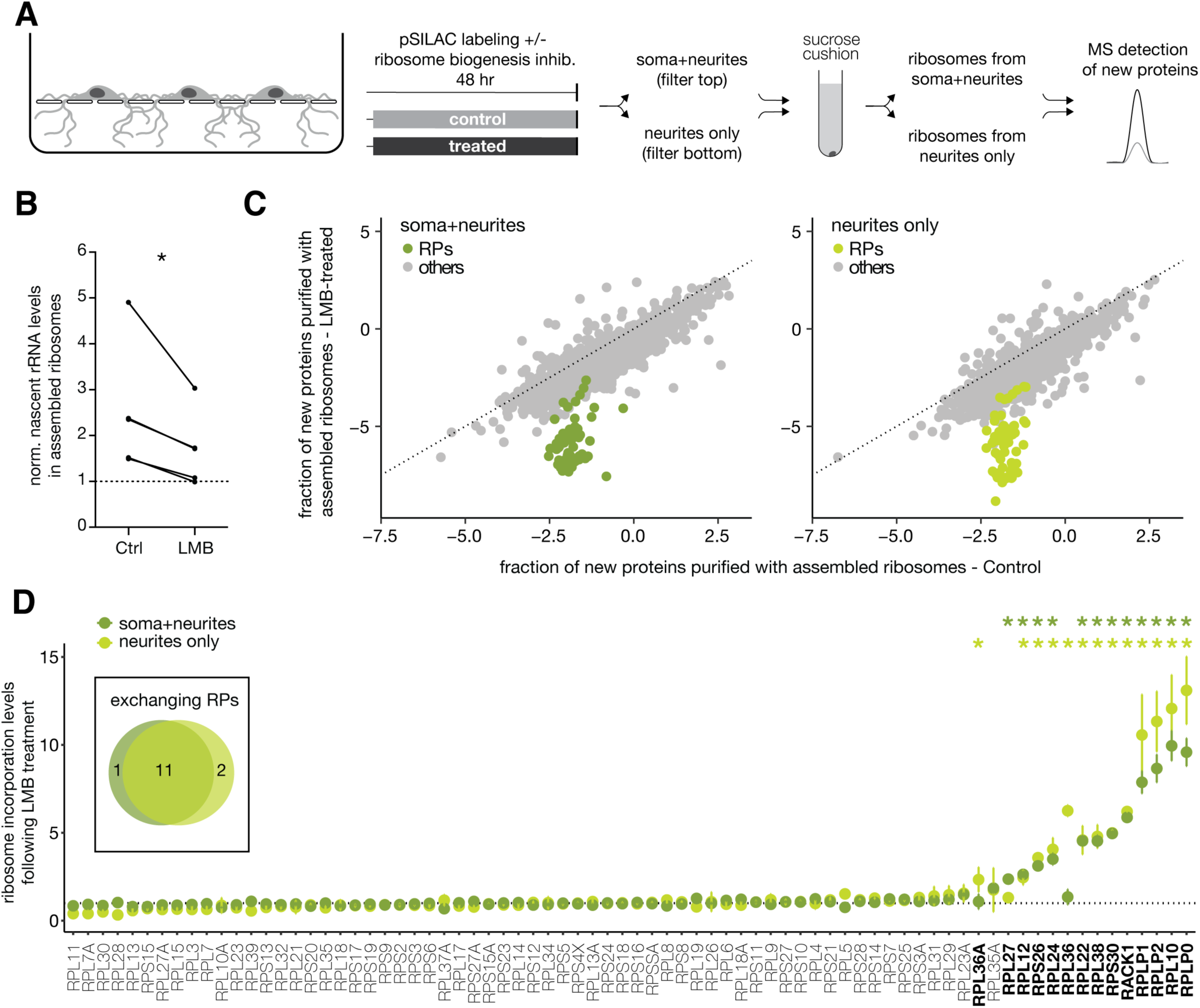
Some nascent RPs can incorporate into neuronal ribosomes independent of the canonical ribosome biogenesis pathway. (A) Schematic of the experimental design. Neurons were cultured on porous membranes where neurites can grow through the pores and be separately harvested. Normal medium was replaced by medium containing heavy amino acids for 2 days (filled boxes) in the presence (treated) or absence (control) of Leptomycin B (LMB) to prevent the export of nascent ribosomes from the nucleus and thus inhibit ribosome biogenesis. Mature ribosomes were purified from either compartment (top: cell bodies + neurites; bottom: neurites only) and new RPs were quantified by mass spectrometry as before. (B) Quantification by RT-PCR of new 18S rRNA in assembled ribosomes, relative to a non-labeled control (y=1, dashed line). 2 days of Leptomycin B (LMB) treatment significantly reduced the nascent rRNA levels in purified ribosomes (5 paired biological replicates, paired *t* test, * p≤0.05). (C) Scatterplots showing fraction of new proteins (log_2_ of the H/L ratio) purified with assembled ribosomes from the cell bodies+neurites (left panel) or neurite-enriched (right panel) compartments of control or LMB-treated neurons. Ribosomal proteins are colored in green, all other proteins are indicated in grey. Dashed lines represent x = y. (D) Fold change of the fraction of new RPs (H/(H+L)) detected in assembled ribosomes between control and LMB-treated samples, normalized to the median value obtained for each subunit (40S or 60S) (see methods). Ribosomes were purified from the cell bodies+neurites (dark green) or neurite-enriched (light green) compartments. Each point represents the mean +/- standard deviation of four biological replicates. Exchanging RPs were identified as significant outliers by ROUT Method (* FDR≤0.5%). Insert: Venn diagram of exchanging RPs in the two compartments.

With LMB as a validated tool to interfere with ribosome biogenesis, we addressed the spatial location of the RP-ribosome dynamics. To do so, we used compartmentalized chambers (Alvarez-Castelao et al., 2020; Poon et al., 2006), which separate and enrich a population of dendrites/axons that can be compared to a mixed cell-body + neurites population that resides above a porous membrane (Figures 4A and S7G-J). We labeled newly synthesized proteins by pSILAC for 48 hrs in the presence or absence of LMB and then purified translating ribosomes from both compartments and measured the fraction of new proteins by mass spectrometry (Figure 4A). LMB treatment did not affect global protein levels in the whole cell lysate (Figure S8A-C), but resulted in a small decrease in total protein synthesis (Figure S8B-D), reproduced also by polysome profiling (Figure S8E-F). Consistent with the average half-life for neuronal ribosomes of ∼8 days (Dörrbaum et al., 2018; Fornasiero et al., 2018; Stoykova et al., 1983), 48 hrs of LMB treatment resulted only in a small decrease of the total assembled ribosomes level (Figure S9A-C), but in a strong and specific reduction of new ribosomes (Figures 4 and S9B-D). In particular, we noted that the large subunit was more affected than the small subunit (Figure S9E). Taken together these data confirmed the successful inhibition of ribosome biogenesis via LMB treatment, prompting us to examine the association of nascent RPs with mature translating ribosomes. We found that while the association of most new RPs was equally reduced by LMB treatment, there was a subset of (∼12) nascent RPs (putative exchangers) that again exhibited a significantly elevated association with translating ribosomes obtained from mixed somata + neurites (dark green data in Figure 4d). As expected, within this group we again identified RPLP1 and 2, which are known to associate with the ribosome in the cytoplasm (Warner and Udem, 1972) and RACK1, which has been recently shown to transiently interact with the ribosome *in vitro* (Johnson et al., 2019). The remainder of the exchanging RPs included RPLP0, RPL10, RPL22, RPL24, RPL38, RPS26, the same RPs that exhibited rapid incorporation in our previous experiment (Figure S10a and Table S2). When we examined the neurite fraction, we found that a largely overlapping group of RPs also showed evidence for nucleus-independent exchange (light green data in Figure 4D). Interestingly, 3 RPs were significantly different in either the mixed (somata + neurites; RPL27) or neurite-enriched compartment (RPL36 and RPL36a) (Figure 4D), suggesting differential exchange of RPs could depend on specific subcellular environments. Additionally, among the common exchangers, several RPs showed a higher incorporation in the neurite compartment, where the contribution of neurites is not diluted by the cell bodies. Taken together, these observations are consistent with the idea that neurite-synthesized RPs can be locally incorporated into pre-existing ribosomes.

The above described exchange of RPs could endow neurons (and other cells) with the ability to repair or remodel ribosomes *in situ* (e.g. (Pulk et al., 2010)) while avoiding the long time delays and the high energetic costs of degrading and producing a whole new ribosome (Granneman and Tollervey, 2007; Warner, 1999). In this regard, we noted a significant negative correlation between the level of incorporation of an RP and its half-life in both cultured neurons and intact brain, with exchanging RPs showing both the shortest and decidedly extreme half-lives when compared to other RPs (Figure S10B-C, (Dörrbaum et al., 2018; Fornasiero et al., 2018)). In addition, the exchanging RPs identified here were present at sub-stoichiometric levels in individual ribosomal subunits quantified in heterologous cells using MS (Imami et al., 2018) (Figure S10D). Furthermore, we evaluated the position of exchanging RPs in the small and large subunits and noted that exchanging RPs were more surface exposed (Figure 5A-B). These data indicate that most of the exchanging RPs detected here differ from other RPs in their half-lives, occupancy levels, and position in the mature ribosome. Finally, among the putative exchangers, 7 RPs (RACK1, RPS30, RPS26, RPLP0, RPL10, RPL24 and RPL36A) were reported in other systems to associate with immature ribosomes during the cytosolic phase of biogenesis, and 5 (RPL38, RPL22, RPL12, RPL27 and RPL36) during the nuclear steps (la Cruz et al., 2015). These data indicate that neurons can exploit the spatial and functional domains of ribosome assembly to dynamically incorporate RPs. Physiological modulation of RPs incorporation

**Figure 5.**
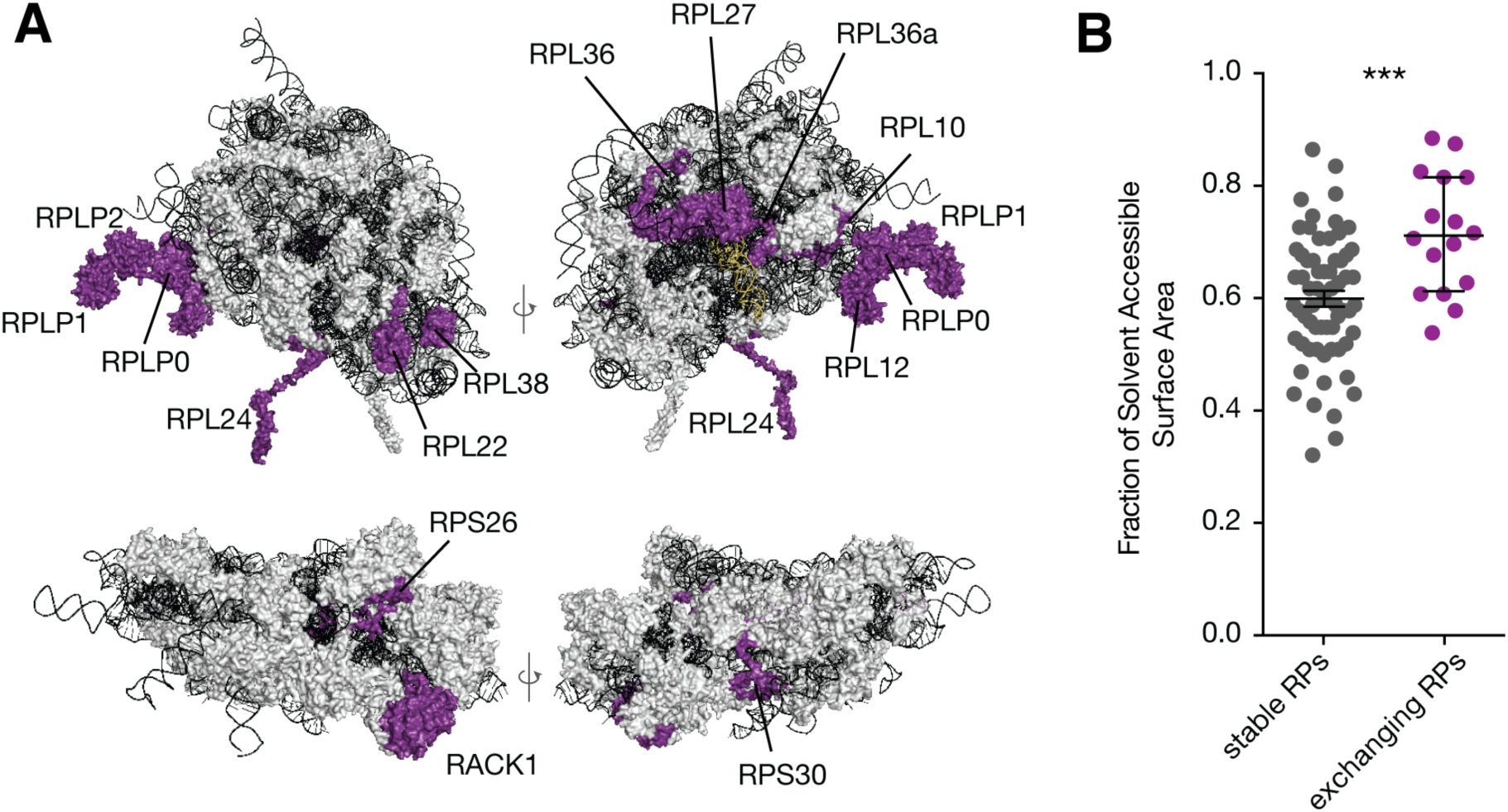
Exchanging RPs tend to be surface-exposed on the assembled ribosome. (A) Structure of human ribosome showing two views each for the large (top) and small (bottom) subunit (PDB: 4v6x). rRNAs in black, tRNA in yellow, stable RPs in grey, exchanging RPs in purple. Names are indicated for the exchanging RPs, as identified in Figure 4D. (B) Fraction of Solvent Accessible Surface Area for stable and exchanging RPs, calculated for individual proteins within the structure of individual subunits of the human ribosome (PDB: 4v6x). Median +/- interquartile range. Mann-Whitney test, *** p≤0.001.

The observation that RP exchange differs in subcellular compartments led us to investigate whether exchange is also regulated by different cellular states. In particular, we examined RP incorporation after a short induction of oxidative stress, via H_2_O_2_ incubation. As ribosomal proteins can be highly oxidized (Mirzaei and Regnier, 2007) and oxidative stress changes neuronal function (Massaad and Klann, 2011), we reasoned that oxidative stress could stimulate ribosome repair. Additionally, oxidative stress rapidly leads to a translation reprogramming, where the synthesis of most proteins is transiently inhibited concomitant with the enhanced translation of specific stress-response transcripts (Kang and Lee, 2001; Shenton et al., 2006; Wu et al., 2019). To rapidly induce oxidative stress, we incubated neurons with H_2_O_2_ for 10 min, during a 3 hrs pSILAC incubation to label newly synthesized proteins (Figure 6A). Consistent with the general inhibition of translation, we found that the overall fraction of new RPs in assembled ribosomes was reduced during stress (Figure S11A). However, the association of a small subset (∼4) of exchanging RPs was relatively enhanced after oxidative stress (Figure 6B-C and Figure S11B-C) while the incorporation of the other exchangers (e.g. RPL24 and RPL22, Figure 6C) was not differentially regulated. Altogether our data indicate that the incorporation probability of different RPs can change according to subcellular environments and physiological conditions.

**Figure 6.**
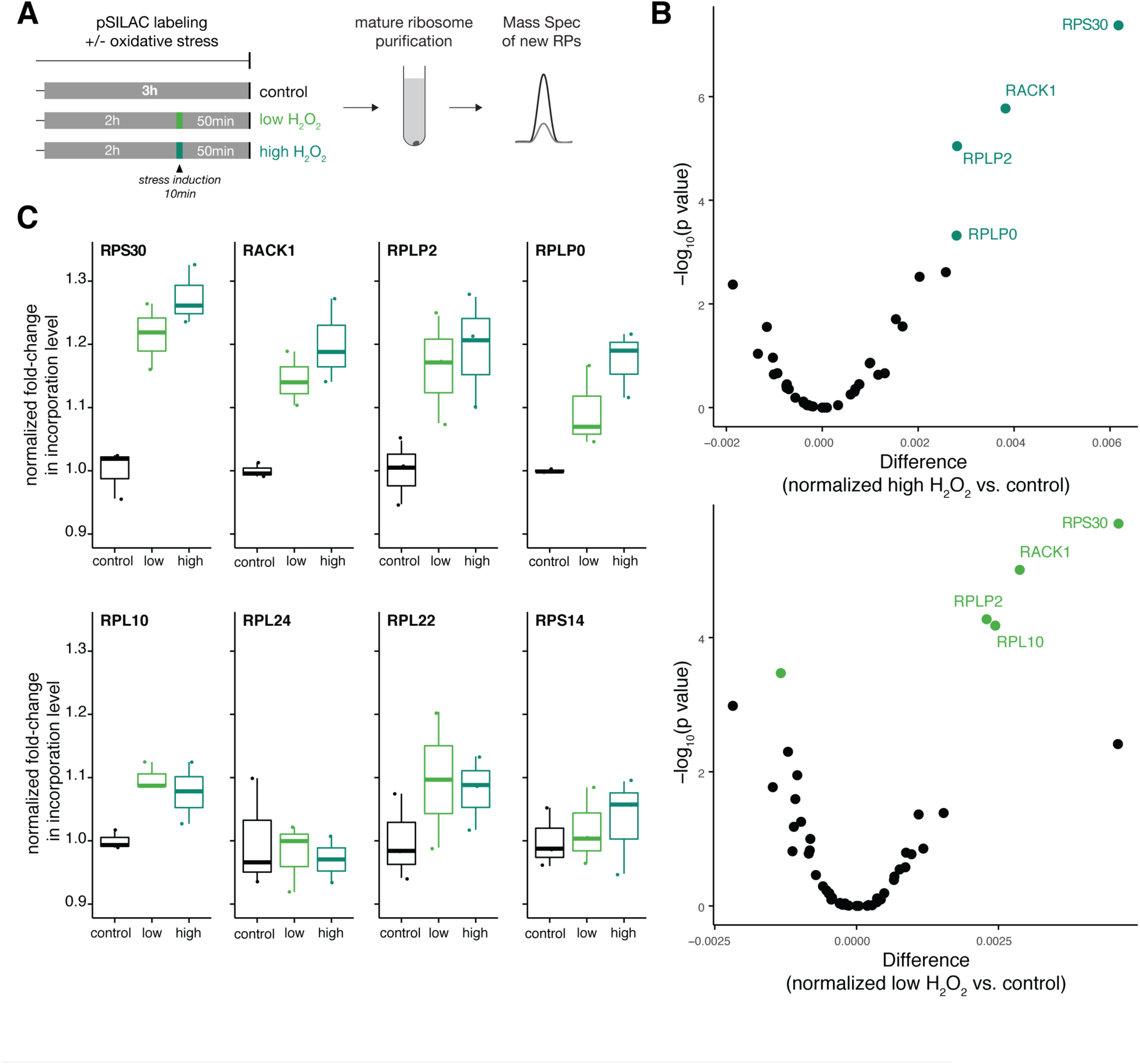
The association of a subset of exchanging RPs is enhanced after oxidative stress. (A) Schematic of the experimental design for measuring new RPs in assembled ribosomes after oxidative stress. Normal medium was replaced by medium containing heavy amino acids (pSILAC) for a total of 3 hr. To induce stress, H_2_O_2_ was added for 10 min to a final concentration of 0.1mM (light green) or 1mM (dark green) to the heavy medium, 2 hrs after the beginning of labeling. Mature ribosomes were purified and new RPs were quantified by mass spectrometry as before. (B) Volcano plots of significantly regulated ribosomal proteins (green, FDR<0.01) in assembled ribosomes, comparing control and H_2_O_2_-treated neurons (see methods). (C) Normalized fold-change in incorporation levels (see methods) for representative ribosomal proteins, which are significantly regulated (RPS30, RACK1, RPLP2, RPLP0 and RPL10) or not (RPL24, RPL22 and RPS14) with H_2_O_2_ treatment. Three biological replicates.

## DISCUSSION

We describe here the localization and translation of ribosomal protein mRNAs in the dendrites and/or axons of neurons. Using pulsed SILAC and a targeted mass spec approach, we measured the incorporation rate of individual nascent RPs into mature ribosomes and identified a subset of ∼12 RPs that exhibited an atypical rapid association with the ribosome. The rapid incorporation of these 12 RPs persisted when ribosome biogenesis was inhibited, providing strong evidence for the exchange of these of RPs on pre-assembled mature ribosomes. Some of the exchanging RPs exhibited an enhanced association with ribosomes in dendrites and axons. Consistent with this, a recent study in *Xenopus* retinal ganglion cells also identified a subset of nascent RPs that associate with ribosomes within axons and the local synthesis of at least one RP was required for axon branching (Shigeoka et al., 2019) (Figure S10E). Interestingly, although we (and others) have observed a large (∼ 50-70) population of RP mRNAs in neuronal processes, under the conditions explored here we detected the dynamic exchange of 12 individual RPs. We note that infrequent incorporation events of new RPs with slower kinetics might be undetectable by our method which involves extremely brief metabolic labeling and labor-intensive manual curation of individual RP nascent peptides. Additionally, under different physiological conditions (e.g. development, stress or synaptic plasticity), some latent localized RP mRNAs could be translated and associate with mature ribosomes. In this case, the large number of distally localized RP mRNAs represents a huge potential for differential RPs incorporation.

Our discovery of dynamic non-canonical incorporation of RPs into mature ribosomes suggests new regulatory scenarios for local translational control. In neurons, the capacity for remote remodeling or repair of ribosomes could be particularly advantageous considering the enormous cell volume, most of which arises from dendrites and axons. In addition, as ribosomes are the most heavily oxidized class of proteins (Mirzaei and Regnier, 2007) and dendrites are particularly sensitive to oxidative insults (Grimm et al., 2018), ribosomes in neuronal processes may be prone to higher proteotoxic damage, creating demand for the local repair of ribosomes. Indeed, our data indicate that oxidative stress promotes the exchange of some RPs on mature ribosomes.

The dynamic incorporation of RPs could also alter ribosome composition resulting in potentially “specialized” ribosomes. The rather long life of the ribosome as a macromolecular protein-RNA machine as well as its nucleocentric biogenesis represent challenges for rapid remodeling. In other systems, ribosomes with altered stoichiometries of RACK1, RPL38 or RPS26 preferentially translate different subsets of mRNAs (Ferretti et al., 2017; Majzoub et al., 2014; Xue et al., 2015). Such specialization could be rapidly achieved through the dynamic exchange of these proteins. This property could be particularly exploited by ribosomes in dendrites and axons which are optimally positioned to respond to synaptic signals that could, in principle, remodel local ribosomes as well as the local translatome.

## Acknowledgements

We thank M. Jovanovic for valuable advice and feedback, A. Biever, C. Glock and G. Tushev for assistance with the translatome data, I. Bartnik, N. Fuerst and C. Thum for the preparation of primary cell cultures, F. Rupprecht for MS maintenance and assistance with data acquisition, C. Polisseni for assistance with image acquisition, M. Heumueller for help with image analysis, I. Epstein for pilot experiments, and M. Beck for comments on the manuscript.

## Author Contributions

Conceptualization: CMF, EMS

Investigation: CMF, KD, ARD, MW, AS, ICC, EV, VV

Formal analysis: CMF, KD, ARD, JDL

Visualization: CMF, KD

Supervision: EMS

Writing – original draft: CMF, EMS

Writing – review & editing: CMF, KD, ARD, MW, AS, ICC, EV, VV, JDL, EMS

## Data and materials availability

All data are available in the main text or the supplementary materials. Analysis scripts are available upon request. The mass spectrometry proteomics data have been deposited to the ProteomeXchange Consortium via the PRIDE (Pérez-Riverol et al., 2019) partner repository with the dataset identifier PXD024678.

---

**Figure S1:**
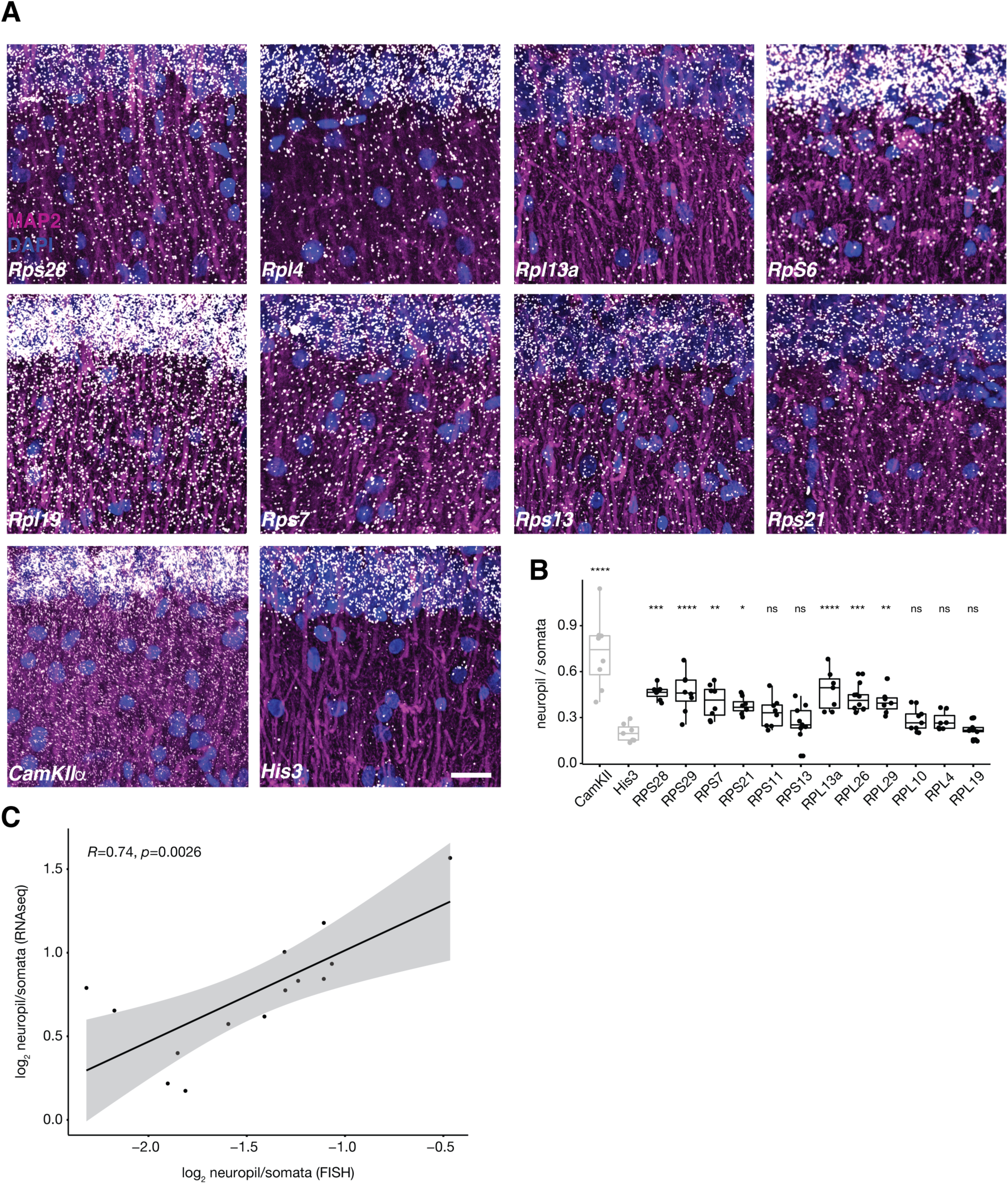
Detection of RP mRNAs in dendrites of rat hippocampal slices. (A) FISH detection of indicated RP mRNAs in dendrites (magenta MAP2, white FISH, blue DAPI) in hippocampal slices. Images are oriented with the somata layer at the top and the dendrites extending below. Scale bar = 50 µm. (B) Quantification of the FISH signal between the neuropil and somata layers in hippocampal slices (relative to the His3 levels, Wilcox test, * p≤0.05, ** p≤0.01, *** p≤0.001, **** p≤0.0001). (C) Pearson correlation plot of the log_2_ expression of the neuropil : somata ratios of mRNA levels measured by FISH and by RNAseq (data sourced from (Biever et al., 2020)).

**Figure S2:**
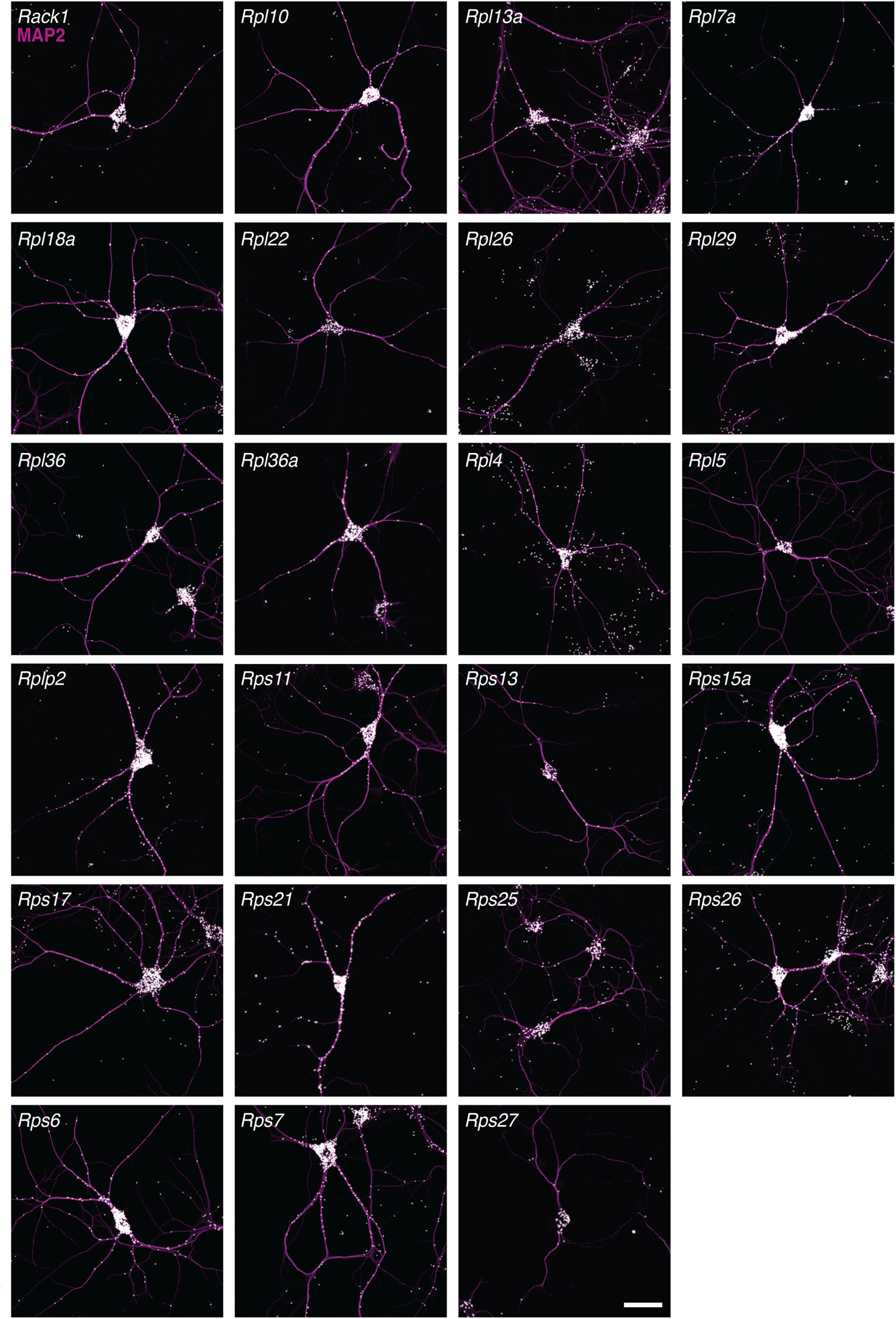
Detection of RP mRNAs in the dendrites of cultured hippocampal neurons. FISH detection of indicated RP mRNAs (magenta MAP2, white FISH) in cultured hippocampal neurons Scale bar = 50 µm. Analysis for these data is shown in Figure 1F.

**Figure S3:**
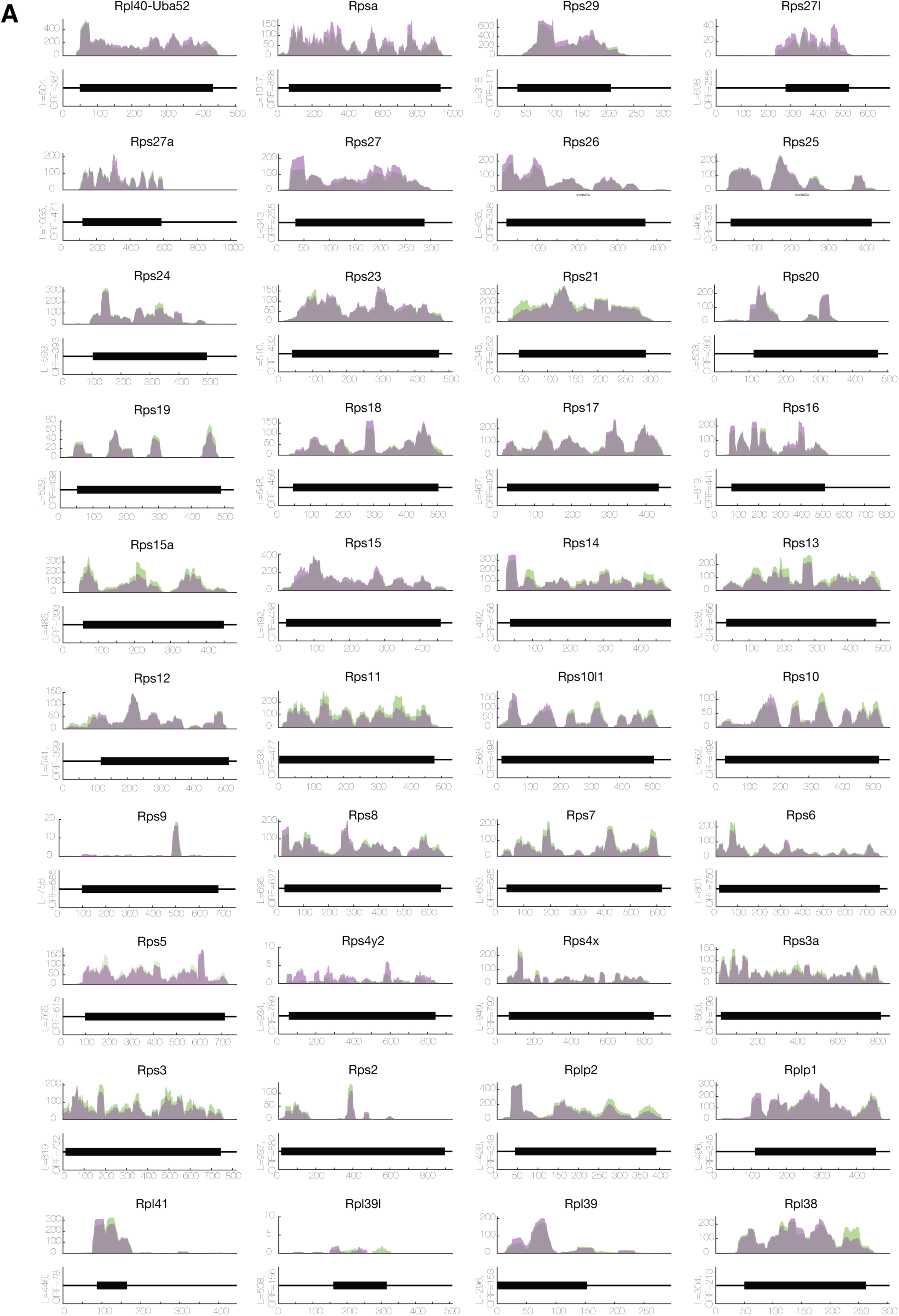
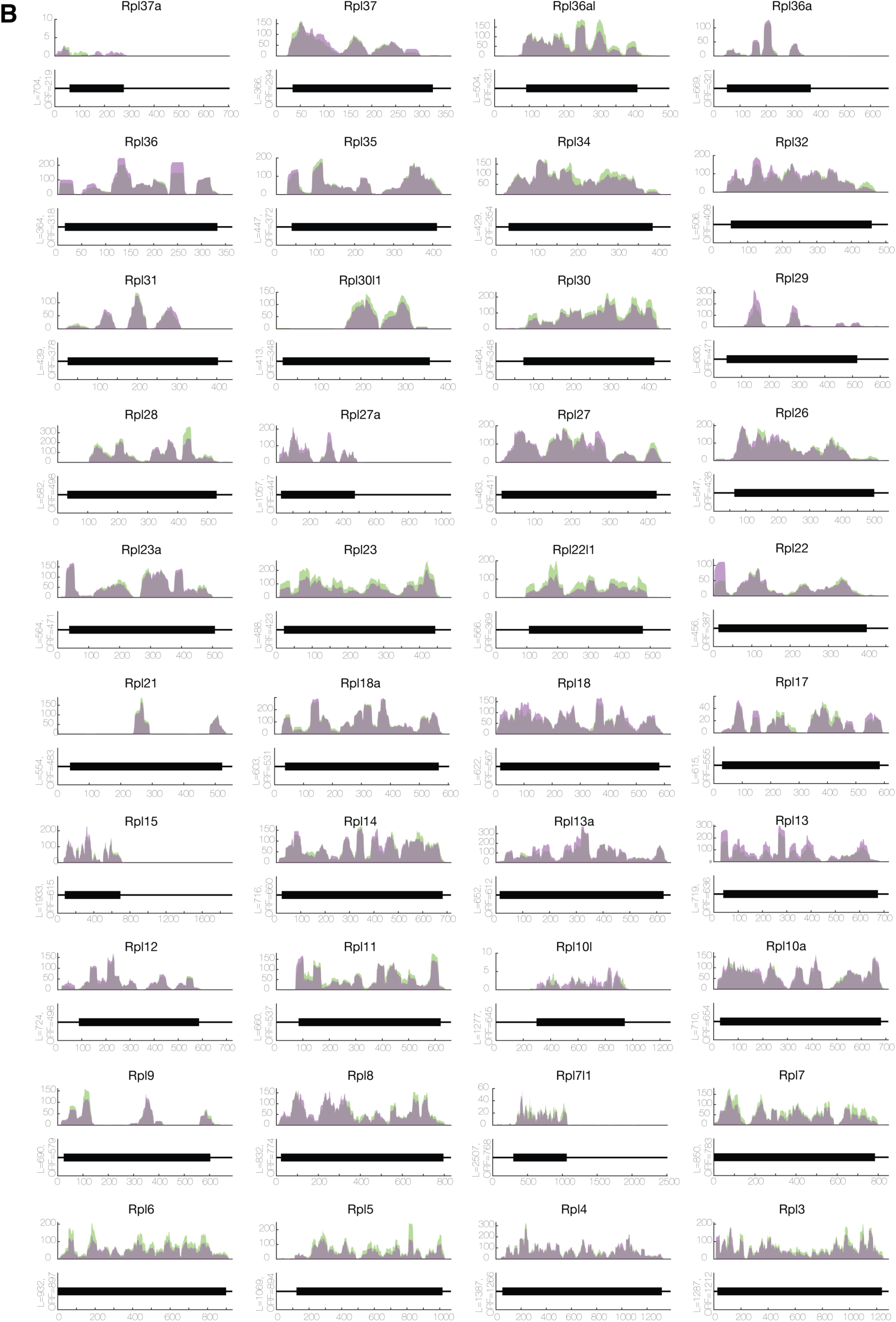
Ribosome footprint coverage of RP mRNAs in rat hippocampal slices. (A-B) Ribosome footprint coverage of RP mRNAs from the somata and neuropil of hippocampal slices (Biever et al., 2020). Shown are the number of reads throughout the open reading frame (black box) from the somata-enriched fraction (green) or the neuropil-enriched fraction (purple). Regions of overlap appear as grey.

**Figure S4:**
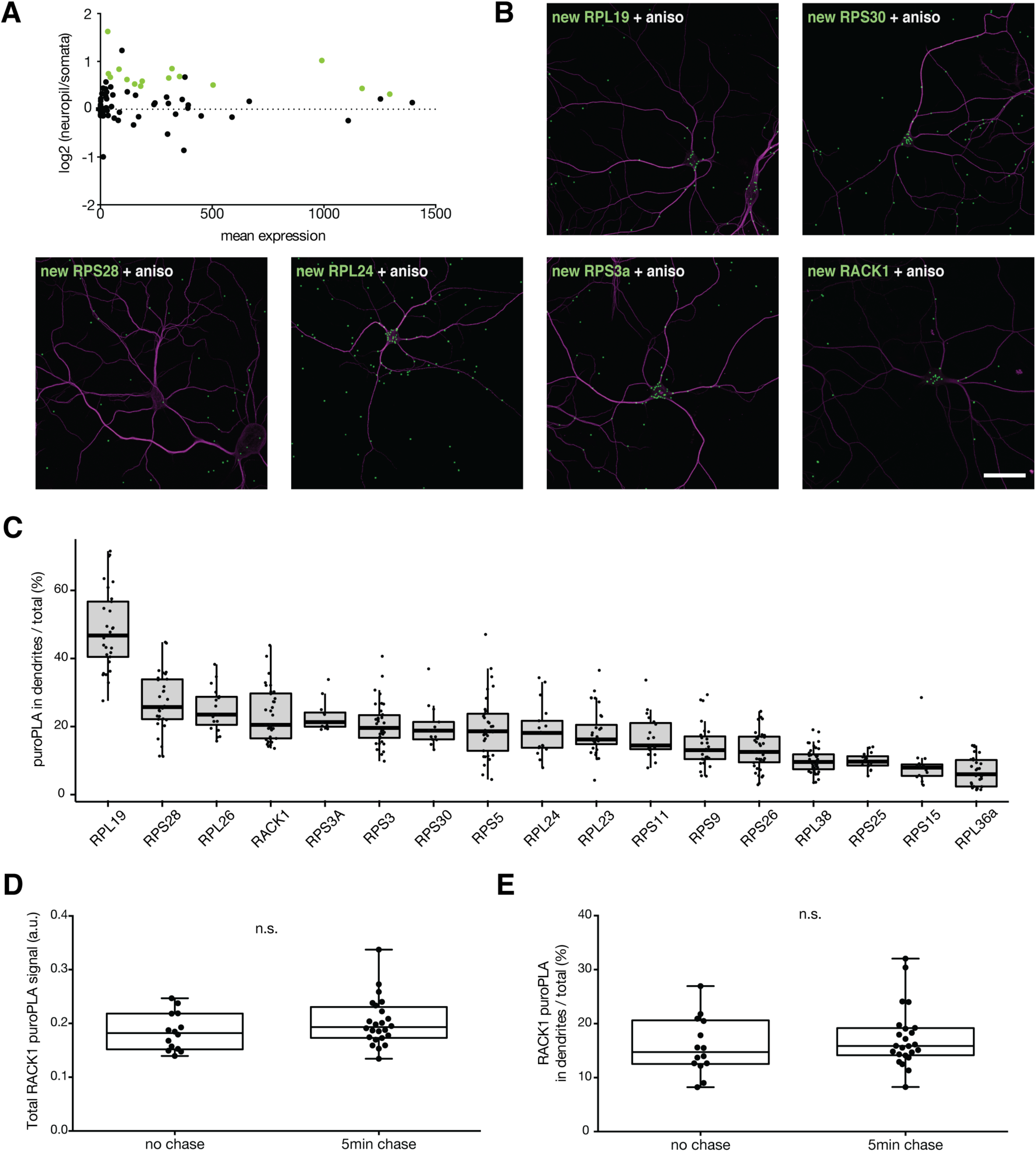
Quantification of dendritic RP mRNA translation. (A) Log_2_ expression of the neuropil : somata ratios of RP footprints (data sourced from (Biever et al., 2020)). RP mRNAs with a significant (adjusted p-value < 0.05) translation increase between the two compartments are highlighted in green. No transcript exhibited a somata-enhanced translation whereas 15 RP mRNAs showed a neuropil-enhanced translation. (B) Puro-PLA detection of nascent RPs in dendrites (magenta MAP2, green newly synthesized RP) in the presence of protein synthesis inhibitor anisomycin (compare to the images shown in Figure 2B). Scale bar = 50 µm. (C) Percentage of nascent RP signal in dendrites over total detected in individual neurons. Proteins are ranked according to their mean values. (D-E) Total levels of nascent RACK1 in a neuron (D) or percentage of the signal in dendrites (E) immediately (no chase) or 5 min after (chase) labeling. Unpaired *t* test, n.s. p>0.5.

**Figure S5:**
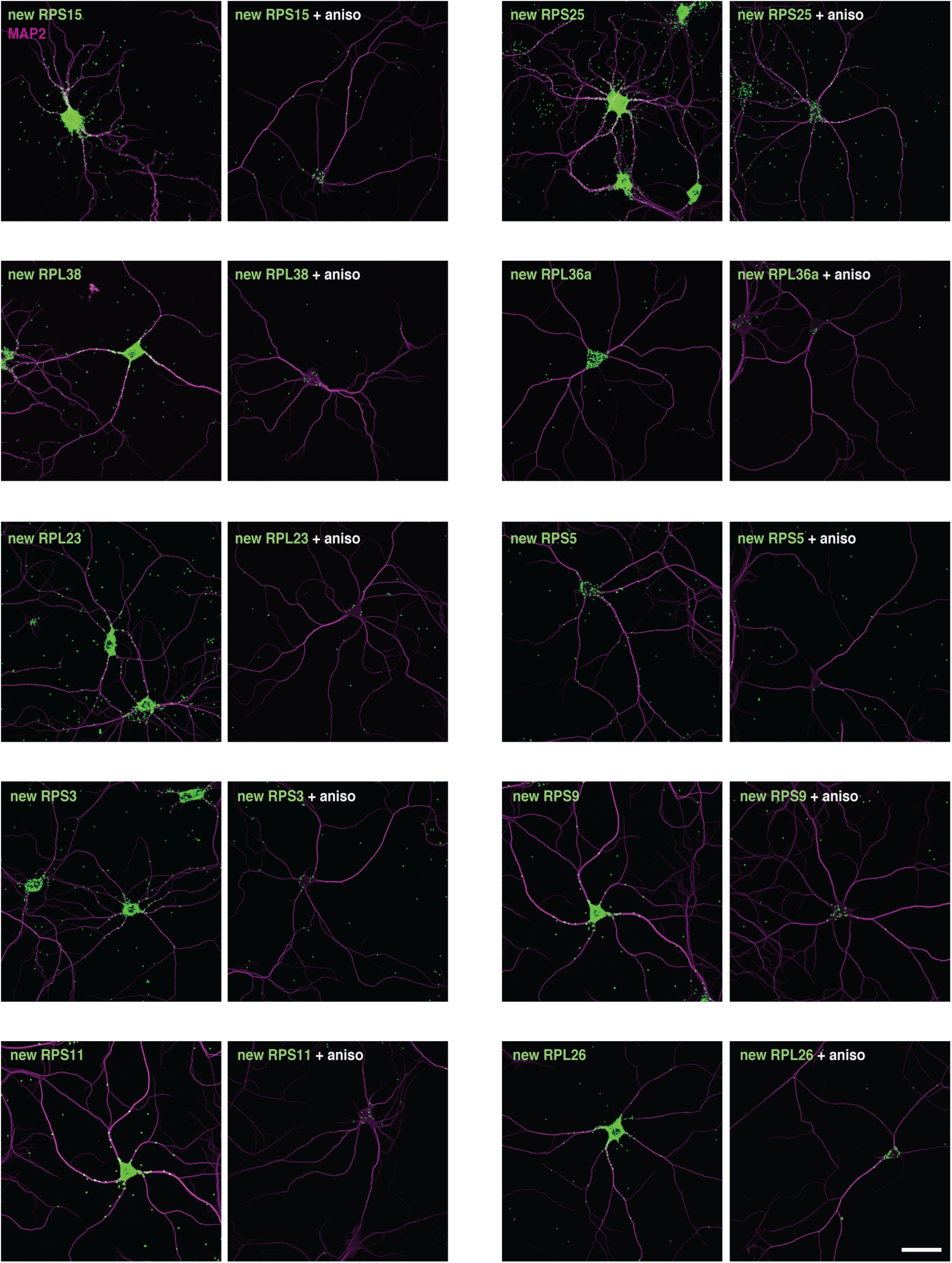
Detection of nascent RPs in dendrites of cultured hippocampal neurons. Detection of nascent RPs (green) in dendrites (magenta for MAP2 immunostaining) of cultured hippocampal neurons using Puro-PLA (tom Dieck et al., 2015). Nascent proteins were labeled with puromycin (5 min), in the absence or presence (as indicated) of the protein synthesis inhibitor anisomycin (see methods). Scale bar = 50 µm.

**Figure S6:**
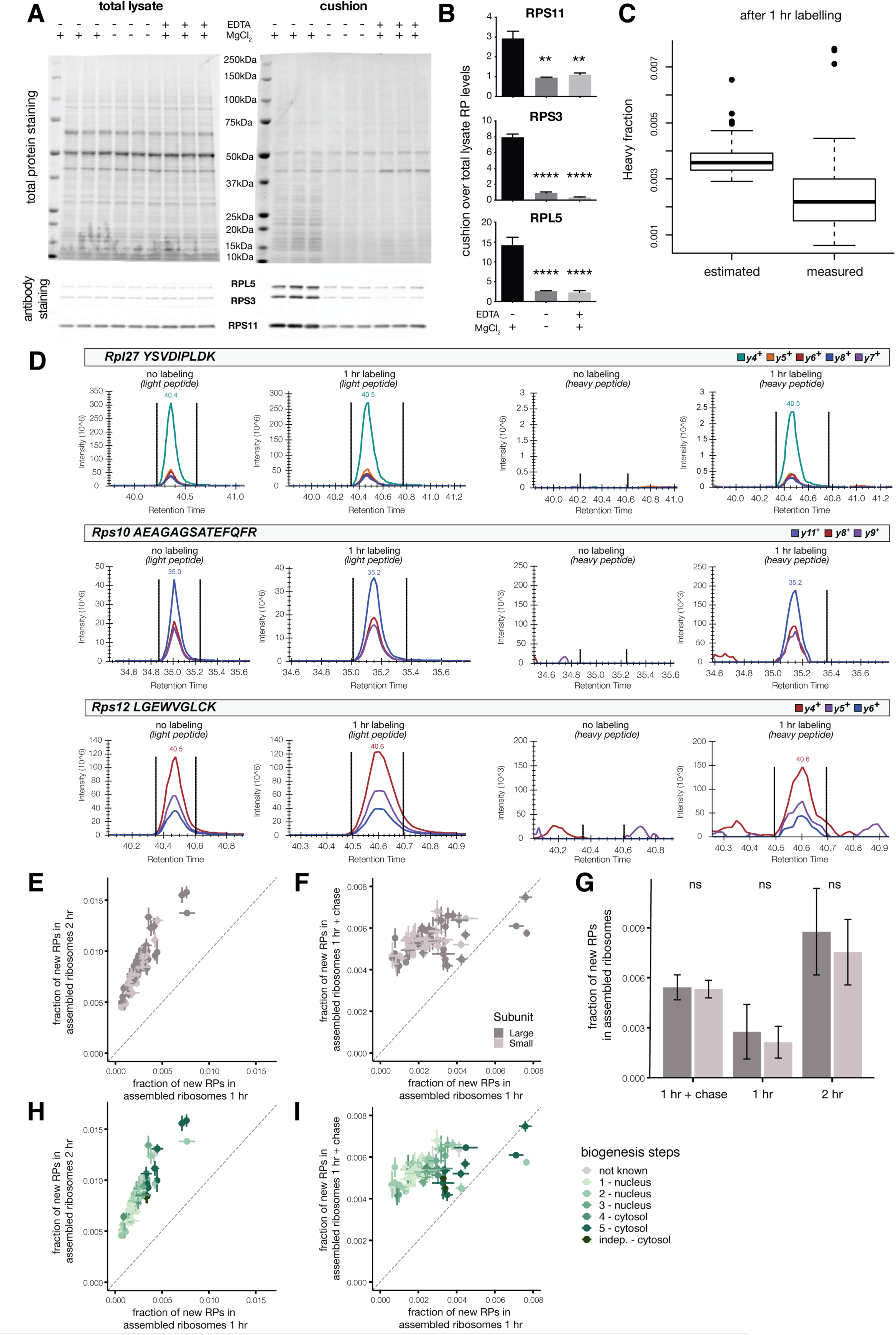
Quality controls for ribosome purification and mass spec detection of newly synthesized RPs. (A) Total protein and Western blot analysis of total lysates (left panel) or cushion samples (right panel) from three biological replicates. Cushion was performed under control conditions or in the absence of magnesium or in the presence of the chelating agent EDTA, as indicated. First lane contains molecular weight ladder. The same percentage of volumes was loaded for lysates and for cushion samples. Total protein (top panel) or RPs of interest (bottom panel) were visualized. (B) Quantification of Figure S6A. The presence of individual RPs was significantly impaired when the cushion was performed under conditions that disassemble monosomes and polysomes into free small and large subunits. Three biological replicates. One-way ANOVA, with Dunnett’s multiple comparisons test. (C) Fraction of labeled peptides (H/(H+L)) for ribosomal proteins, after 1 hr of labeling with heavy amino acids, estimated based on the half-life quantified by Dörrbaum et al. (Dörrbaum et al., 2018) or measured in our data set. (D) Examples of the traces for heavy or light peptides measured by Mass Spectrometry with Parallel Reaction Monitoring (see methods), after 1 hr of labeling or not, as indicated. Representative peptides from the indicated RPs of high (top panel), medium (middle panel) or low (bottom panel) abundance were chosen. (E-F) Related to Figure 3C-D. Scatterplots showing fraction of new RPs (H/(H+L)) in assembled ribosomes after different labeling conditions, as indicated in x- and y- axes. Points represent average +/- standard deviation of three biological replicates. Proteins are colored according to which ribosomal subunit they belong to (small subunit in light gray, large subunit in dark gray). Dashed line represents x = y. (G) Related to Figure 3E-F. Mean +/- standard deviation of the fraction of new proteins (H/(H+L)) in assembled ribosomes for RPs of the same subunit (small subunit in light gray, large subunit in dark gray). Different labeling conditions are indicated on the x-axis. Wilcox test, n.s. p>0.05 (H-I) Related to Figure 3C-D. Scatterplots showing fraction of new RPs (H/(H+L)) in assembled ribosomes after different labeling conditions, as indicated by the x- and y- axes. Points represent average +/- standard deviation of three biological replicates. Proteins are colored according to which step they are known to incorporate into immature ribosomes during biogenesis (as reviewed in (la Cruz et al., 2015)). The darkest green color represents RPLP1 and RPLP2, whose incorporation is not linked to ribosome biogenesis. Dashed line represents x = y.

**Figure S7:**
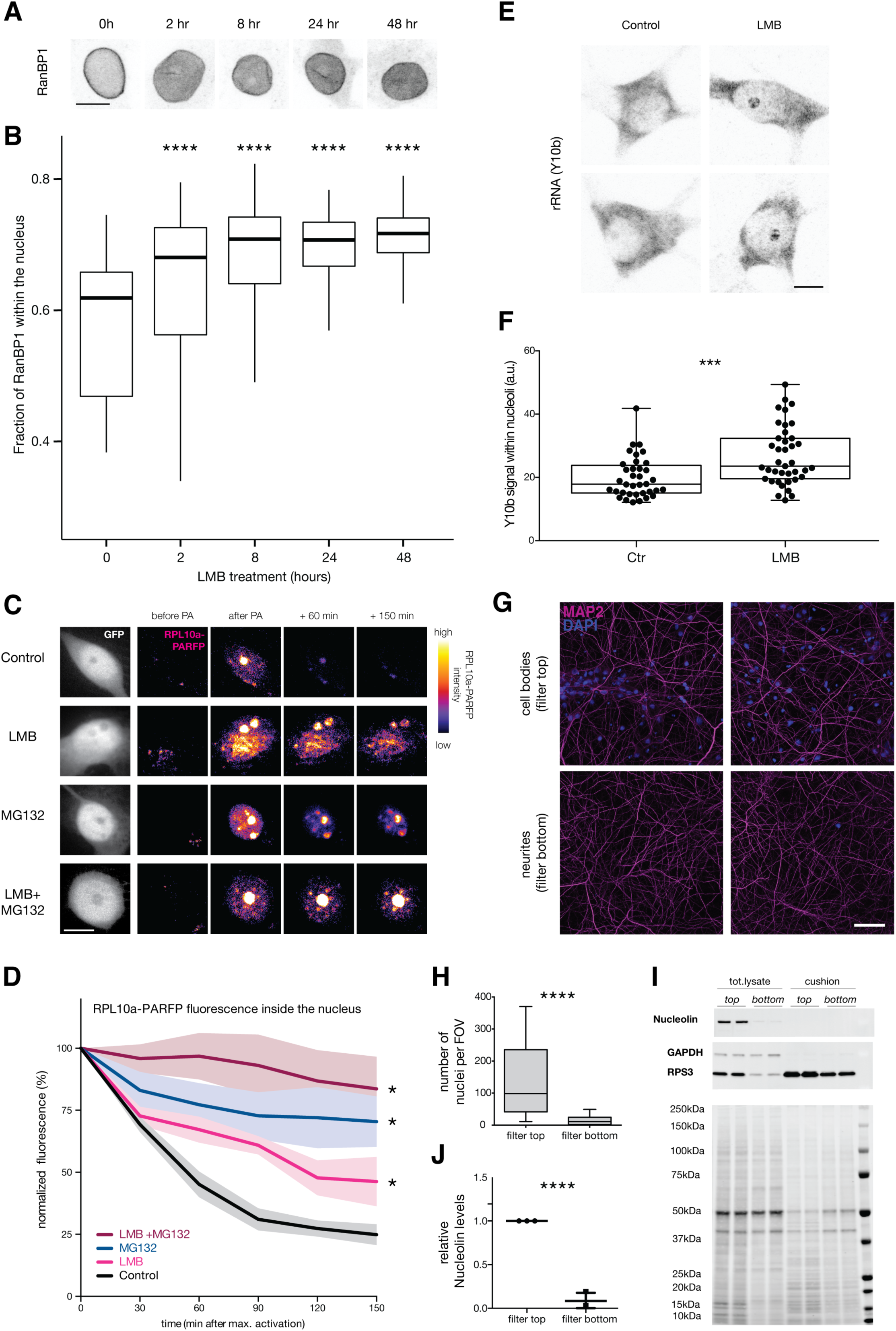
Leptomycin B (LMB) rapidly blocks CMR1-mediated nuclear export, and thus sequesters nascent ribosomes in the nucleus. Compartmentalized chambers allow for the enrichment of neurites. (A) Immunolabeling of RanBP1 in neuronal cell bodies at different durations of LMB incubation. RanBP1 is an accessory protein involved in the CRM1-mediated transport, known for its rapid shuffling between nucleus and cytoplasm. Scale bar = 10µm. (B) The fraction of RanBP1 signal within the nucleus significantly increases during LMB incubation at all time points tested (Wilcox test, p < 0.001). (C) Representative images of neurons transfected with GFP (grey) and RPL10a-photo-activatable (PA)-RFP (fire color look-up), before and after photoactivation, under control conditions (first row) or in the presence of LMB to block nuclear export (second row), or MG132 to block proteasome-mediated degradation (third row), or both LMB and MG132 (fourth row). Scale bar = 10 µm. (D) Percentage of RPL10a-PARFP fluorescence inside the nucleus, normalized to the maximum nuclear intensity reached after photo-activation. Both the individual and combined inhibition of nuclear export and proteasomal degradation significantly slowed down the decay of RPL10a-PARFP signal in the nucleus (mean +/- SEM, repeated measures ANOVA, p < 0.01, Tukey’s multiple comparisons test, p <0.05). (E) Immunolabeling of rRNA (Y10b antibody) in neuronal cell bodies under control condition or after 2 days of LMB treatment. Scale bar = 10µm. (F) The intensity of Y10b signal inside nucleoli significantly increases after 2 days of LMB incubation (unpaired *t* test, p<0.001) (G) Representative images indicating the presence of dendrites (MAP2) and nuclei (DAPI) in the compartmentalized chamber. Top or bottom view of the filter, as indicated (scale bar 50 µm). (H) Analysis of nuclear de-enrichment in the compartmentalized chambers. The number of nuclei per Field Of View was dramatically decreased in neurite compartment (bottom). Five biological replicates, for a total of > 45 Fields of View per compartment. Unpaired *t* test, **** p≤0.0001. (I) Total protein and Western blot analysis of lysates or cushion samples from somata + neurites (top) or neurites (bottom) compartments, as indicated. Last lane contains molecular weight ladder. Total protein (bottom panel) or proteins of interest (bottom panel) were visualized: Nucleolin as a nuclear marker, GAPDH as a cytosolic marker, RPS3 as a ribosome marker. (J) Quantification of protein analysis. The level of Nucleolin detected in the somata + neurites (top) fraction was significantly higher than that observed in the neurite (bottom) fraction. Three biological replicates. Unpaired *t* test, **** p≤0.0001.

**Figure S8:**
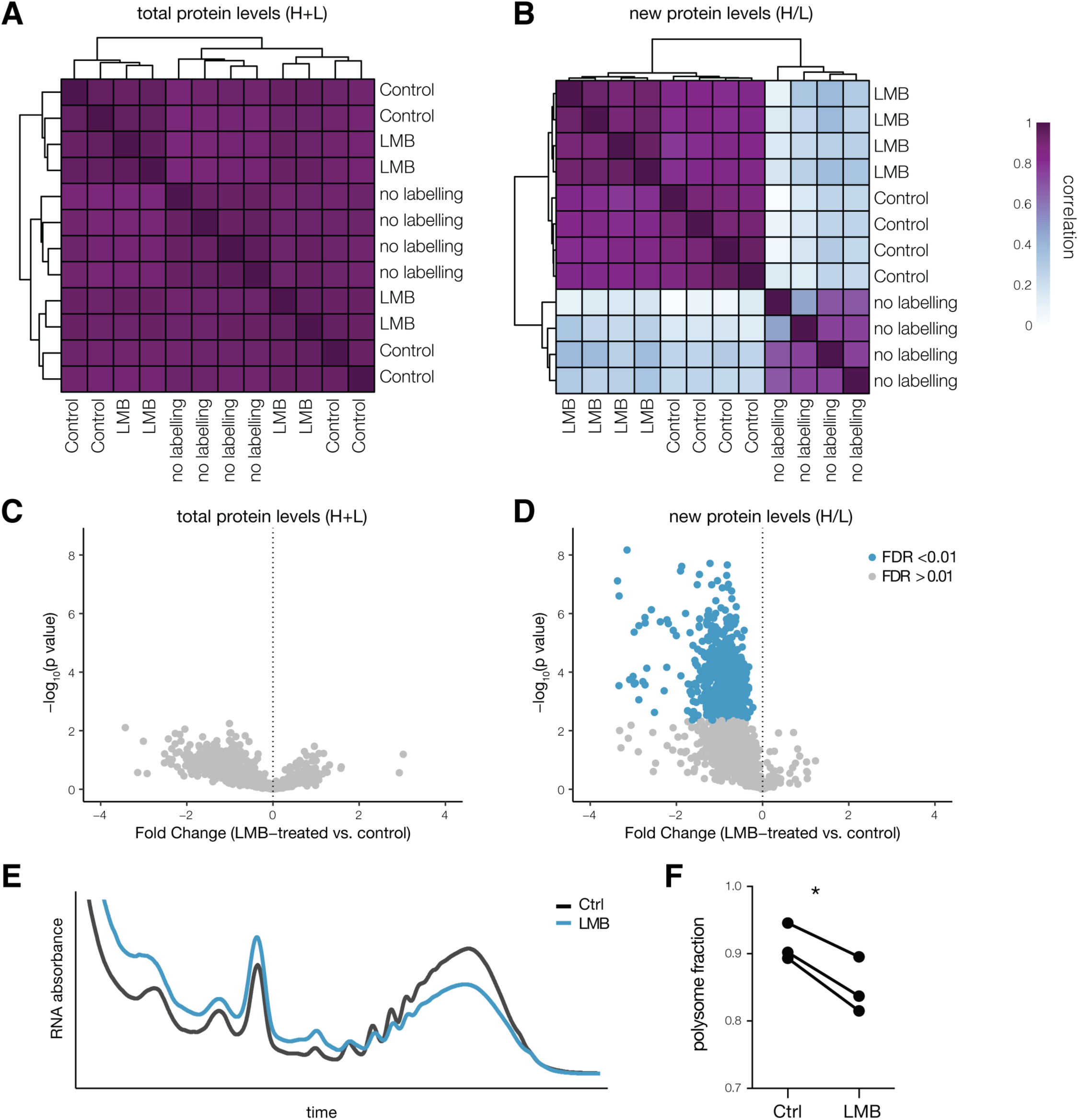
LMB treatment does not affect the total cellular protein levels but does reduce a population of new proteins. (A-B) Hierarchical clustering of biological replicates (see methods) according to the similarity of total (log_2_ of H+L, (A) or new (log_2_ of the H/L ratio, (B) protein levels in lysates from control, LMB-treated or unlabeled neurons. Cells are color coded according to pairwise Pearson correlations. Biological replicates of the same labeling condition cluster together only when comparing the fraction of newly synthesized proteins. (C-D) Volcano plots of significantly regulated proteins (blue, FDR<0.01) in total lysates, comparing control and LMB-treated neurons. Nuclear export inhibition via LMB does not affect total protein levels (log_2_ of H+L, (C) but results in a general decrease in newly synthesized proteins (D). Dashed line represents Fold Change = 0. (E) Representative polysome profiles from control (black) or LMB-treated (blue) neurons. (F) Quantification of the polysome fraction from control and LMB-treated neurons. Ribosome biogenesis inhibition via LMB resulted in a decrease in protein synthesis. Paired *t* test, p<0.05.

**Figure S9:**
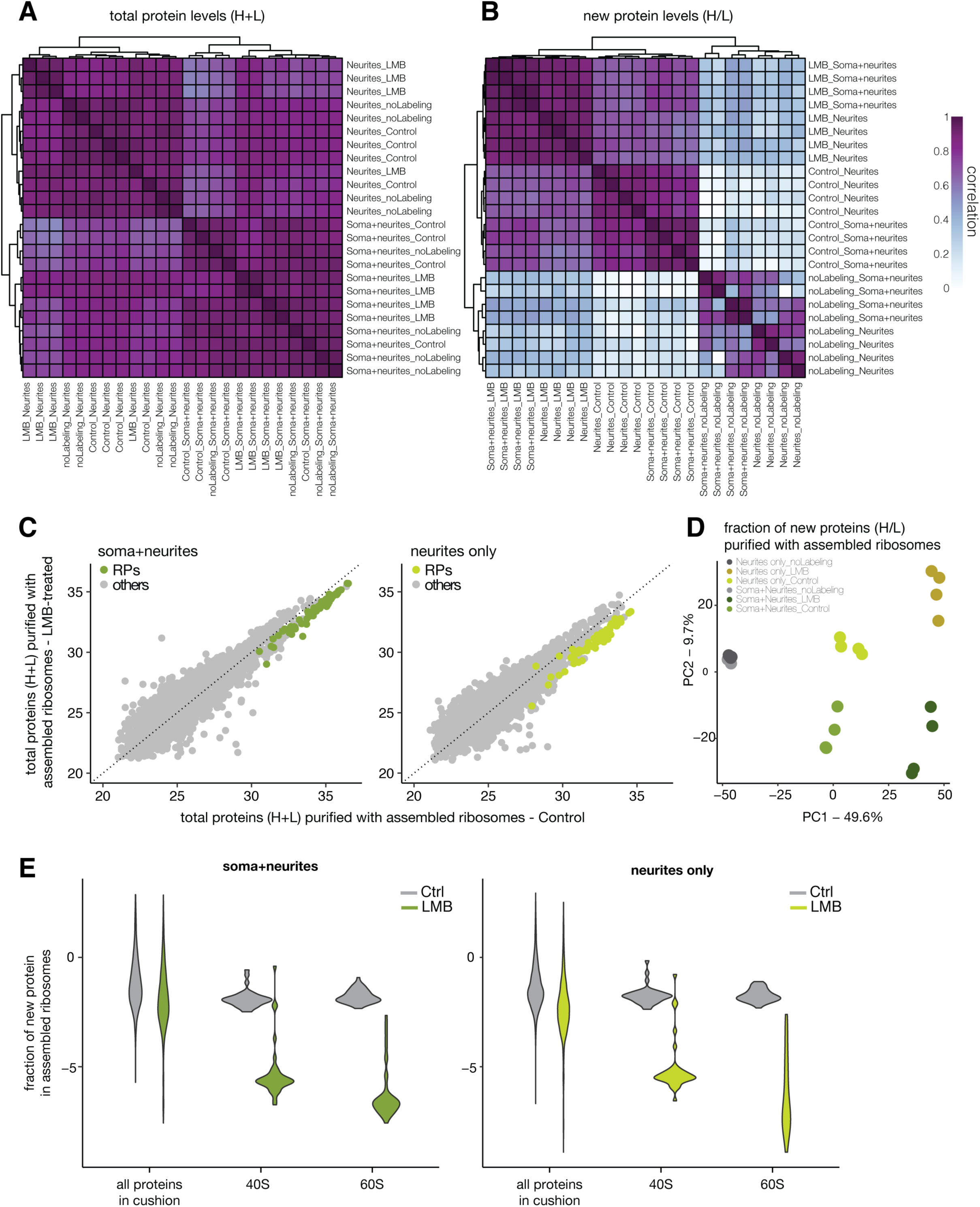
LMB treatment results in less new ribosomes. (A-B) Hierarchical clustering of biological replicates (see methods) according to similarity of total (log_2_ of H+L, (A) or new (log_2_ of the H/L ratio, (B) levels of proteins co-purified with assembled ribosomes from cell bodies or neurites of control, LMB-treated or unlabeled neurons. Cells are color coded according to pairwise Pearson correlations. When comparing total protein levels, the unsupervised clustering segregates samples according to compartment but not according to labeling condition (A). When comparing the fraction of newly synthesized proteins instead, biological replicates of the same labeling condition and from the same compartment are successfully clustered together (B). (C) Scatterplots showing total levels of protein (log_2_ of H+L) co-purified with assembled ribosomes from cell bodies (left panel) or neurites (right panel) of control or LMB-treated neurons. Ribosomal proteins are indicated in green. Average of four biological replicates. Dashed line represents x = y. (D) PCA analysis showing similarities across new protein levels co-purified with assembled ribosomes from cell bodies or neurites of control, LMB-treated or unlabeled neurons. (E) Levels of new proteins (log_2_ of the H/L ratio) co-purified with assembled ribosomes from cell bodies (left panel) or neurites (right panel) of control (grey) or LMB-treated (green) neurons. Ribosomal proteins from either small (40S) or large (60S) subunit are grouped as indicated in the x-axis. Average of four biological replicates.

**Figure S10:**
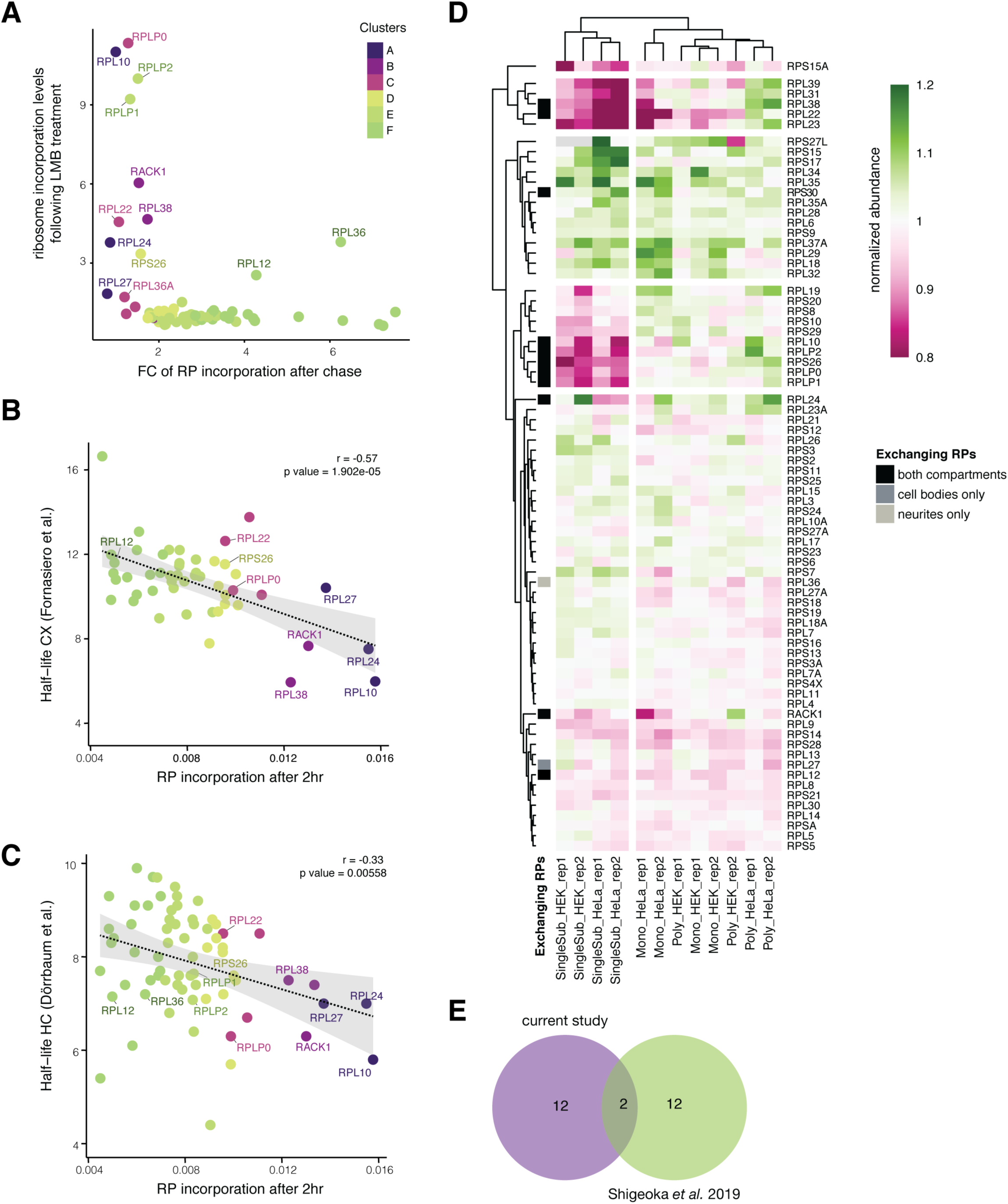
Exchanging RPs tend to be short-lived and sub-stoichiometric in single subunits. (A) Scatterplot showing the fold change of RP incorporation after a chase (x-axis, relative to Figure 3d) and the incorporation levels after LMB treatment (y-axis, average between compartments from Figure 4d). Ribosomal proteins are colored according to the clusters identified in Figure 3b. Exchanging RPs are indicated by name. (B-C) Scatterplots showing the incorporation of new RPs after 2 hrs of labeling (x-axis, as in Figure 3) and their half-life (y-axis) as measured in intact cortical tissue (Fornasiero et al., 2018), (B) or in cultured hippocampal neurons (Dörrbaum et al., 2018), (C) Proteins are colored according to the clusters identified in Figure 3b. Exchanging RPs are indicated by name. Dashed line represents the regression line and the confidence interval is shown in grey. (D) Heatmap showing RP expression levels across different ribosomal conformations (selected fractions within polysome profiling gradients) in HEK and HeLa cells, as measured using MS by (Imami et al., 2018). Pseudocells (protein levels normalized to the median of the corresponding ribosomal subunit) are ordered using unsupervised clustering, both for columns (biological replicate of each condition) and rows (individual ribosomal protein). The exchanging RPs in both compartments are shown in black, those exchanging only in the compartment with cell bodies are shown in dark grey, and those exchanging only in neurites are shown in light grey (as identified in Figure 4d). SingleSub = single ribosomal subunits (large or small). Mono = monosome. Poly = polysome. (E) Venn diagram showing the overlap between the exchanging RPs identified in the current study (purple) and the nascent RPs detected in axonal ribosomes after removal of the cell body in a recent study ((Shigeoka et al., 2019), green). Curiously, the two known positive controls of exchange, RPLP1 and P2, were not detected in (Shigeoka et al., 2019).

**Figure S11:**
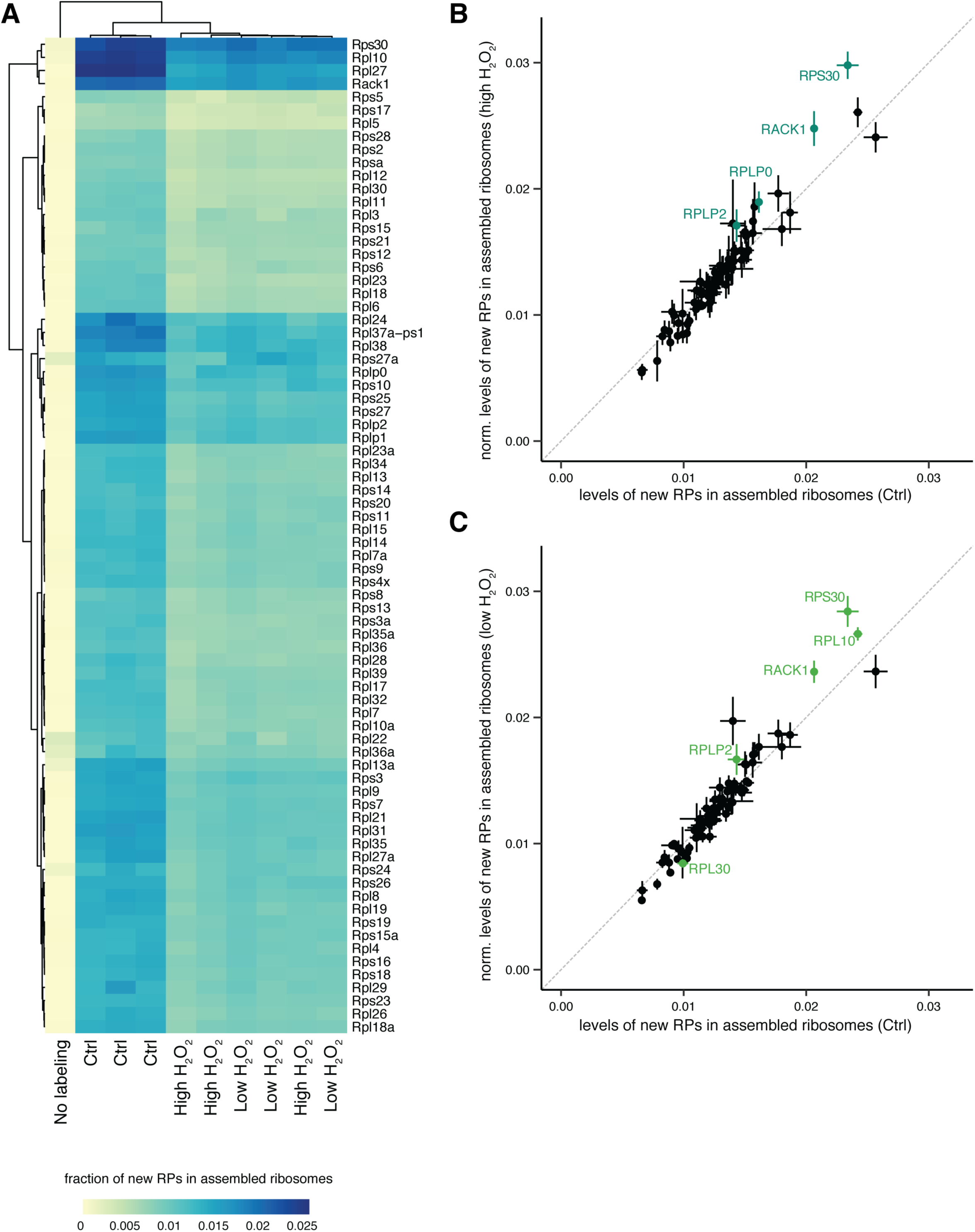
Effect of oxidative stress on RPs incorporation into assembled ribosomes. (A) Heatmap showing for each RP the fraction of new proteins (H/(H+L)) incorporated into assembled ribosomes. Pseudocells (median of peptides obtained per individual protein) are ordered according to unsupervised clustering, both for columns (biological replicate of each condition) and rows (individual ribosomal protein). Experimental conditions of the labeling are indicated at the bottom. (B-C) Scatterplots showing the fraction of new RPs (H/(H+L)) in assembled ribosomes after the different labeling conditions, as indicated by x- and y- axes. To correct for the general decrease in protein synthesis with stress, the values in the H_2_O_2_ samples were normalized over the mean fold change. Points represent average +/- standard deviation of three biological replicates. Proteins are colored according to the significance (green, FDR<0.01) calculated as in Figure 5B. Dashed line represents x=y.

**Table S1:** Detection of Ribosomal Proteins mRNAs in previously reported studies (related to Figure 1).

**Table S2:** Incorporation levels of nascent Ribosomal Proteins into assembled ribosomes after short labeling (related to Figure 3) and after LMB treatment (related to Figure 4).

**Table S3:** Incorporation levels of nascent Ribosomal Proteins into assembled ribosomes after oxidative stress (related to Figure 5).

**Table S4:** Results of the statistical analysis for the differential incorporation of Ribosomal Proteins between control and high concentration of H_2_O_2_ (related to Figure 5).

**Table S5:** Results of the statistical analysis for the differential incorporation of Ribosomal Proteins between control and low concentration of H_2_O_2_ (related to Figure 5).

**Table S6:** detailed summary of the parameters used for LC+MS.

**Table S7:** detailed summary of the settings used for MaxQuant.

## STAR METHODS

### EXPERIMENTAL MODEL AND SUBJECT DETAILS

#### Preparation of primary cultured neurons

Dissociated rat hippocampal or cortical neuron cultures were prepared and maintained as described previously (Aakalu et al., 2001). Briefly, we dissected hippocampi or cortices from postnatal day 0 to 1 rat pups of either sex (Sprague-Dawley strain; Charles River Laboratories) and dissociated the samples with papain (Sigma). For imaging experiments, hippocampal neurons were plated at a density of 30 × 10^3^ cells/cm^2^ on poly-Dlysine–coated glass-bottom petri dishes (MatTek). For biochemical experiments, cortical neurons were plated at a density of 4 × 10^6^ cells/cm^2^ on poly-Dlysine–coated 10 cm dishes, at a density of 1 × 10^6^ cells/cm^2^ on poly-Dlysine–coated 6 cm dishes or at a density of 9 × 10^6^ cells/cm^2^ on poly-Dlysine–coated 75 mm inserts (3.0 µm pore size, Corning 3420). One day after plating on the inserts, cells were incubated with 5 µM AraC (Sigma C1768) for two days, then the AraC was removed by changing the media. Neurons were maintained in a humidified atmosphere at 37°C and 5% CO_2_ in growth medium [Neurobasal-A supplemented with B27 and GlutaMAX-I (Life Technologies)] for 14-16 DIV for biochemical experiments or for 25-27 DIV for imaging experiments. All experiments complied with national animal care guidelines and the guidelines issued by the Max Planck Society and were approved by local authorities.

#### Pharmacological treatments

To inhibit ribosome biogenesis, cells were treated with 20 µg/µL Leptomycin B (Merck, 431050) for 2 days, unless otherwise specified. To inhibit proteasome degradation, cells were treated with 10 µM MG132 (Sigma, M8699) for at least 2 hr. To induce oxidative stress, cells were incubated with 1mM or 0.1mM H_2_O_2_ (AlfaAesar L13235) for 10 min.

#### Metabolic labeling of newly synthesized proteins by pSILAC

Heavy medium was prepared by adding 84 mg/L of Arg10 (ThermoFisher 88434) and 146 mg/L of Lys8 (ThermoFisher 88432) into a Arg- and Lys-free Neurobasal-A medium (ThermoFisher, customized). To condition the medium, extra plates from each prep were incubated with the same medium starting from DIV 0. On the day of the pulsed-SILAC experiment (DIV 13-16), the conditioned heavy medium was collected from the extra plates. The original “light” medium was then replaced by the conditioned heavy medium for the indicated amount of time (1 hr, 2 hrs, 3 hrs or 2 days). To reduce the likelihood of purifying polypeptide chains still emerging from the ribosome, a 5 min wash with light medium was used to allow termination of translation after the 1 hr, 2 hrs or 3 hrs labeling.

### METHOD DETAILS

#### Ribosome purification by sucrose cushioning

Cells were washed three times and scraped in ice-cold DPBS (ThermoFisher, 14040-091) supplemented with 100 µg/mL of CHX (Sigma, C7698). An aliquot was saved to prepare a whole cell lysate. Cells were pelleted by 5 min centrifugation at 500 x *g*. For ribosome purification, cells were lysed in 400 µL of ribosome lysis buffer (20 mM Tris pH 7.4, 150 mM NaCl, 5 mM MgCl2, 24 U/mL TurboDNase, 100 µg/mL cycloheximide, 1% TritonX-100, 1 mM DTT, RNasin(R) Plus RNase inhibitor 200U/mL and 1x cOmplete EDTA-free protease inhibitor). For the compartmentalized chambers, first the top compartment (cell bodies + neurites) was scraped in DPBS+CHX and then the bottom (neurites only) was scraped directly in 200 µL of ribosome lysis buffer. To guarantee sufficient yield from the neurite compartment, each biological replicate consisted of the content of two inserts pooled together. Lysates were pipetted up and down until homogenization was clear with a 0.4×20mm syringe needle (HSW FINE-JECT) on ice. Samples were then centrifuged at 10,000 x *g* for 10 min at 4°C. Supernatants were loaded on 1 mL sucrose solution (34% sucrose, 20 mM Tris pH 7.4, 150 mM NaCl, 5 mM MgCl2, 1 mM DTT, 100 µg/mL cycloheximide) in a thickwall polycarbonate tube (Beckman, 349622) and centrifuged for 30 min at 4°C at 55000 rpm with a SW55Ti rotor (acceleration 0, deceleration 7). Ribosome pellet was resuspended in 20 µL of 10 mM HEPES, 120 mM NaCl, 3 mM KCl, 10 mM D-Glucose, 2mM MgSO4 and 2 mM MgSO4, and submitted to Mass Spectrometry or Western Blot Analysis. For the experiment shown in Figure S6a-b, ribosome lysis buffers and cushion solutions were modified to either contain 0 mM MgCl2 or 5mM MgCl2 + 15 mM EDTA.

#### Ribosome purification by polysome profiling

Cells were processed as for sucrose cushioning, except the ribosome lysis buffer was supplemented with 8% glycerol and the supernatant were loaded on a 10-50% sucrose gradient. For the gradients, all solutions were prepared in gradient buffer (20 mM Tris pH 7.5, 8% glycerol, 150 mM NaCl, 5 mM MgCl2, 100 µg/mL cycloheximide, 1 mM DTT). Gradients were prepared by sequentially adding solutions with different sucrose concentrations (in order from first added to last, 8 mL of 55%, 0.5 mL of 50%, 0.5 mL of 40%, 0.5 mL of 30%, 0.5 mL of 20%, 0.5 mL of 10%) into the same Thinwall polypropylene tube (Beckman, 331372). Tubes were placed at −80°C to freeze the content before adding the next sucrose solution, and finally stored at −80°C. The day prior to experiments, gradients were left for equilibration at 4°C overnight. Then 1 to 2 OD (measured with NanoDrop at 260 nm) of the lysates were loaded on top of the gradients and spun at 36,000 rpm at 4°C for 2 hrs with a SW41-Ti rotor (Beckman). Gradients were then run at 850 µL/min in a density gradient fractionation system (Teledyne Isco), chased by 60% sucrose 10% glycerol in water. RNA absorbance at 254 nm was continuously measured using a UA-6 detector. The area under the curve corresponding to the monosome and polysomes was measured. To compare across different runs, the polysome fraction was calculated as area under polysomes over the sum of the areas under monosome and polysomes.

#### Total cell lysates

Cell were scraped as described above for sucrose cushioning. Cell pellets were lysed in 200 µL of 8 M urea, 200 mM Tris/HCl [pH 8.4], 4% CHAPS, 1 M NaCl, cOmplete EDTA-free protease inhibitor (Roche, 11873580001), using a pestle. Lysates were sonicated at 4°C for 4 rounds of 30 sec each, and incubated for 10 min with 1 µL of Benzonase (Sigma E1014). After centrifugation for 5 min at 10,000 x *g*, the supernatant was submitted to Mass Spectrometry or Western Blot validation.

#### FISH in hippocampal slices

3-5 weeks-old Sprague Dawley SPF rats were housed on a 12/12-hour light/dark cycle with food and water ad libitum until euthanasia. Animals were anesthetized by Isoflurane inhalation (Abbott, USA). The rat head was removed and immediately frozen in liquid nitrogen for 30 s. The brain was extracted and sliced into 500-600 µm slices in ice-cold oxygenated sucrose-ACSF using a vibratome (VT1200S, Leica, Germany). The slices were fixed in a fixation buffer (4 % PFA, 4 % sucrose in PBS) for 1 hr at 4 °C and then 1 hr at room temperature. After washing with PBS, slices were dehydrated with ice-cold 15 % sucrose in PBS for 2-3 hrs at 4 °C and then ice-cold 30 % sucrose in PBS overnight at 4 °C. Slices were blocked in O.C.T. (SAKURA Finetek USA Inc., USA) and sliced again at 30 µm thickness using a sliding microtome (Microm HM450, ThermoFisher), followed by thorough washing with PBS and fixation for 20 min at room temperature using a fixation buffer (4 % paraformaldehyde, 5.4 % glucose, 0.01 M sodium metaperiodate in lysine-phosphate buffer). *In situ* hybridization was performed using the ViewRNA ISH Cell Assay Kit (ThermoFisher) according to the manufacturer’s instructions with some modifications. Briefly, slices were permeabilized at room temperature using the detergent solution for 20 min. After washing with PBS and 5 min incubation with the hybridization buffer, the respective probes (see table below for details) were diluted 1:100 in the pre-warmed working hybridization buffer and added to the slices. After incubation at 40 °C overnight and washing with the wash buffer, 1:100 PreAmplifier Mix was diluted in pre-warmed working amplifier diluent and incubated with the slices for 1 hr at 40 °C, followed by 1 hr incubation with 1:100 Amplifier mix in pre-warmed working amplifier diluent and then 1 hr with 1:100 Label probe mix in pre-warmed working label probe diluent. After washing, the slices were permeabilized with 0.5 % Triton X-100 in blocking buffer (4 % goat serum in PBS) for 20 min and blocked in blocking buffer for 1 hr. Immunostaining was then performed using antibodies against Map2 (see table below for details) overnight at 4 °C, followed by secondary antibody donkey anti-gp Cy5 (1:500, 706-175-148, Dianova, Germany) and 1:1000 DAPI for 2 hrs at room temperature. Mounting of slices was performed using Aqua Poly/mount (18606, Polysciences, USA).

#### FISH in hippocampal cultures

Cultured rat hippocampal neurons (DIV 21-28) were fixed for 20 min at room temperature using a fixation solution (4 % paraformaldehyde, 5.4 % glucose, 0.01 M sodium metaperiodate in lysine-phosphate buffer). *In situ* hybridization was performed using the ViewRNA ISH Cell Assay Kit (ThermoFisher) according to the manufacturer’s instructions with some modifications. Briefly, neurons were permeabilized at room temperature by treating with detergent solution for 5 min, followed by pepsin digestion (0.01 mg/ml of enzyme in 10 mM HCl) for 45 sec. After washing with PBS, the respective probes (see table below for details) were diluted 1:100 in pre-warmed hybridization buffer and added to neurons. After incubation at 40 °C for 3 hr, neurons were washed with wash buffer and stored in storage buffer overnight at 4 °C. After several washes, neurons were incubated for 30 min at 40 °C with PreAmplifier Mix (diluted 1:25 in pre-warmed working amplifier diluent), followed by 30 min incubation with 1:25 Amplifier mix in pre-warmed working amplifier diluent and then 30 min with 1:25 Label probe mix in pre-warmed working label probe diluent. After washes, neurons were immunostained with antibodies against Map2 (see table below for details) overnight at 4 °C, followed by secondary antibody donkey anti-gp Cy5 (1:500, 706-175-148, Dianova, Germany) for 1 hr at room temperature and DAPI for 5 min.

#### Puro-PLA

Detection of newly synthesized proteins by puromycin labeling and proximity ligation was performed as previously described (tom Dieck et al., 2015). Neurons were metabolically labeled for 5 min with 1 µM puromycin (Sigma, P8833). For the chase experiment (shown in Figure S3d-e), after the 5 min labeling cells were washed three times and incubated for 5 more minutes with the original medium. All samples were washed twice with DPBS (ThermoFisher, 14040-091) prior to fixation (20min in 4% PFA in 4% sucrose in PBS). Cells were permeabilized (15 min in blocking buffer + 0.5% Triton-X 100) and blocked (>30 min in blocking buffer, PBS + 4% goat serum). Neurons were incubated overnight at 4°C in PBS + 4% goat serum containing primary antibodies against puromycin, the protein-of-interest and MAP2 to label dendrites (see table below for details). After washing, Proximity Ligation Assay (PLA) was performed using the Duolink In Situ PLA kit (Sigma). In particular, PLA probes anti-rabbit PLUS (DUO92002) and anti-mouse MINUS (DUO92004) and the Duolink Detection reagents Red (Sigma DUO92008) were used according to the manufacturer’s recommendations. Briefly, probes (1:10 dilution) and a secondary antibody for MAP2 were applied in PBS with 4% goat serum for 1 h at 37 °C, washed three times with wash buffer A (0.01 M Tris, 0.15 M NaCl, 0.05% Tween 20) and incubated for 30 min at 37°C with the ligation reaction. Samples were then washed three times with wash buffer A and incubated at 37 °C for 100 min with the amplification reaction mixture. Amplification was stopped by three washes in wash buffer B (0.2 M Tris, 0.1 M NaCl, pH 7.5). Nuclei were stained with DAPI (1:1000 for 2 min) and cells were kept in wash buffer B at 4°C until imaging.

#### Image acquisition and analysis for Puro-PLA and FISH

Within a week after labeling, samples were imaged using a LSM780 confocal microscopy (Zeiss) and a Plan-Apochromat 40x/1.4 Oil DIC M27 objective. Z-stack was set to cover the entire volume of a neuron, with optical slice thickness set to optimal. Laser power and detector gain were adjusted to avoid saturated pixels. Imaging conditions were held constant within experiments. Maximum intensity projections of image z-stacks were used for image analysis. For visualization purposes (but not for analysis), the punctae size was dilated and brightness and contrast adjusted. Image analysis was performed in ImageJ/FIJI with an in-house script. For cell culture, the dendritic arbor and the cell body of individual neurons were manually traced using the MAP2 immunolabeling. For hippocampal slices, the somatic compartment was defined by a 5 µm dilation of the DAPI signal. After thresholding, the intensity and number of punctae were quantified and normalized over the annotated neuronal area.

#### Immunofluorescence

Cells were fixed for 20min in 4% PFA in 4% sucrose in PBS, permeabilized for 15 min in 0.5% Triton-X 100 + blocking buffer and blocked for at least 30 min in blocking buffer (PBS + 4% goat serum). Neurons were incubated for 2 hrs with primary antibodies and, after three washes, for 1 hr with secondary antibodies, all in blocking buffer (see table below for antibodies information). After two washes in PBS, cells were stained with DAPI (1:1000 for 2 min) and kept in PBS at 4°C until imaging. For validation of the compartmentalized chambers, pieces of the filter were processed as described above, and mounted on a glass slide (ThermoFisher 10417002) with Aqua Poly/mount (Polysciences, 18606). Samples were imaged using a LSM780 confocal microscopy (Zeiss) using a Plan-Apochromat 40x/1.4 Oil DIC M27 or Plan-Apochromat 20x/0.8 M27 objectives. A Z-stack was set to cover the entire volume of neurons, with optical slice thickness set to optimal. Laser power and detector gain were adjusted to avoid saturated pixels. Imaging conditions were held constant within experiments. Maximum intensity projections of z-stacks were used for image analysis. For visualization purposes (but not analyses) brightness and contrast were adjusted.

All image analyses were performed in ImageJ/FIJI with a fully automated script built in-house. In Figure S6a-b, the intensity of RanBP1 signal was quantified within two masks, containing the whole nucleus with or without the outer edge (representing the nuclear envelope). The fraction of RanBP1 within the nucleus was calculated as the signal in the inner mask, over the signal in the outer mask. In Figure S6e-f, the intensity of the Y10b signal was quantified within a mask based on the Nucleolin channel, which was co-stained to label nucleoli. In Figure S6g-h, the number of nuclei was quantified based on the DAPI channel.

#### Mass spectrometry

##### Sample preparation for MS analysis

Proteins were digested according to the ‘Filter-Aided Sample Preparation’ (FASP) protocol (Wiśniewski et al., 2009) or using S-Traps according to an adapted version of the suspension trapping protocol described by the manufacturer (ProtiFi, Huntington, NY). Peptides were desalted using C18 StageTips (Rappsilber et al., 2007), dried by vacuum centrifugation and stored at -20°C until LC-MS analysis.

##### LC-MS/MS Analysis

The peptide samples were reconstituted in 5% acetonitrile (ACN) and 0.1% formic acid (FA) supplemented with an iRT peptide standard (1:10 dilution; Ref.: Ki-3002-2; Biognosys). Peptides were separated by nano-HPLC (U3000 RSLCnano, Dionex). The samples were loaded and washed with loading buffer (2% ACN, 0.05% trifluoroacetic acid (TFA) in water; 6 min; 6 µL/min) on a PepMap100 loading column (C18, L = 20 mm, 3 µm particle size, Thermo Scientific) and subsequently separated on a PepMap RSLC analytical column (C18, L = 50 cm, <2 µm particle size, Thermo Scientific) by a gradient of phase A (0.1% FA in water) and phase B (80% ACN, 0.1% FA in water). The gradient was ramped from 4% B to 48% B in 90 min at a flow rate of 300 nL/min. All solvents were purchased from Fluka in LC-MS grade. Eluting peptides were ionized online using a Nanospray Flex ion source (Thermo Scientific) and analyzed in a Q-Exactive Plus mass spectrometer (Thermo Scientific) (see Table S6 for method details). In brief, for DDA mode, precursor ion spectra were acquired over the mass range 350–1400 m/z (mass resolution (R)= 70 k, AGC target 3×10^6^, maximum injection time (IT) = 60 ms). The top-10 precursor ions were selected for fragmentation (HCD; normalized collision energy = 30) and analyzed in MS2 mode (R = 17.5 k, isolation window = 1.7 Da, AGC target = 2×10^4^, maximum IT = 50 ms). In a parallel reaction monitoring (PRM) approach (Peterson et al., 2012), MS2 scans were acquired (R = 17.5 k, isolation window = 1.7 Da, AGC target = 1×10^5^, maximum IT = 64 ms) according to the scheduled inclusion lists.

##### MS-data processing

For protein identification and relative quantification of the DDA data, MS raw data were analyzed with MaxQuant (version 1.6.2.3 and 1.6.0.1; RRID:SCR_015753) (Cox and Mann, 2008; Tyanova et al., 2016) using customized Andromeda parameters (see Table S6 for LC+MS parameters and Table 7 for MaxQuant settings). For all searches, spectra were matched to a Rattus norvegicus database (reviewed and unreviewed; downloaded from uniprot.org (RRID:SCR_004426)) considering tryptic peptides with up to 2 missed cleavages and to contaminant and decoy databases. Precursor mass tolerance was set to 4.5 ppm and fragment ion tolerance to 20 ppm. Carbamidomethylation of cysteine residues was set as fixed modification and protein-N-terminal acetylation as well as methionine oxidation were set as variable modifications. A false discovery rate (FDR) of 1% was applied at the peptide-spectrum-match (PSM) and protein level. Only proteins identified by at least one unique peptide were retained for downstream analysis. For relative protein quantification, the data was searched with a multiplicity of 2 (light (Lys0, Arg0) and heavy (Lys8, Arg10)) and the LFQ values were computed without normalization.

For the targeted analysis of ribosomal proteins by PRM, raw data was analyzed in Skyline (version 20.1.0.155; RRID: SCR_014080) (MacLean et al., 2010). To obtain information on target peptides, a series of DDA scout runs was measured. Targeted peptides were selected based on uniqueness, no missed cleavages, recurrent occurrence and signal intensity. Given a high degree of sequence similarity amongst RPs, some ribosomal proteins could not be represented by more than one unique peptide. Peptide identity was confirmed using a spectral library generated in Skyline using the results of a MaxQuant search (msms.txt) with a multiplicity of 1 (only light) of the DDA data. In Skyline, a scheduled method was generated using target detection windows of 3 min which was split into three inclusion lists to analyze separate injections measured with PRM methods 1-3. For final data curation, PRM raw data were imported as multiple-injection replicates in Skyline and peak picking was confirmed manually in accordance to retention time, mass accuracy and library matches.

All MS data associated with this manuscript have been uploaded to the PRIDE repository and are available with the dataset identifier PXD024678 (RRID:SCR_003411; (Pérez-Riverol et al., 2019)). Anonymous reviewer access is available upon request.

##### Protein quantification and statistical analyses

For targeted analysis of nascent ribosomal proteins after 1 hr, 2 hrs or 3 hrs of SILAC labeling, heavy and light peptide signals were curated in Skyline, peak areas were exported and heavy peptide fractions (%H=H/(H+L)) were calculated in R. Protein heavy fractions were determined by combining the peptide-specific heavy fractions by their median value. Protein fold changes were calculated as median of the corresponding peptide fold changes. For unsupervised clustering, protein heavy fractions were hierarchically clustered using Euclidean distance. Visualization of the cluster data was done using the pheatmap R-package (RRID: SCR_016418, https://CRAN.R-project.org/package=pheatmap).

For analysis of nascent ribosomal proteins after 2 days of SILAC labeling with or without LMB treatment, peptide signals of ribosomal proteins were manually curated in Skyline to ensure accurate quantification, especially for low abundant heavy peptides. Subsequently heavy peptide fractions were calculated in R. Technical replicates were merged (mean) and only peptides with a heavy fraction that was 3x greater in the SILAC samples (both the LMB treated and untreated condition) compared to the “no labeling” samples were used for down-stream analysis. LMB-treated versus control fold changes of the heavy fractions were calculated on the peptide level and protein fold changes were determined as the median of the corresponding peptide fold changes. To correct for different size effect of LMB-treatment on the two ribosome subunits, the fold change of each ribosomal protein was normalized over the median change of the corresponding subunit.

To analyze the effect after 2 days of SILAC labeling with or without LMB treatment on all proteins in the total lysate- or cushion-samples, MaxQuant results of the protein groups (proteinGroups.txt) were further processed in R. Protein groups were filtered to remove contaminant or decoy database hits and proteins only identified by a modified peptide (“identified only by site”). Total intensity (H+L) and heavy over light ratios (H/L) were log2-transformed and values of the technical duplicates were merged by their means. Subsequent principal component and Pearson correlation analyses were conducted in R using its base functions. Differential regulation comparing LMB-treated and control samples was investigated using unpaired, two-sided t-tests. To correct for multiple testing, Benjamini-Hochberg correction was applied with an FDR cut-off < 0.01.

For the analysis on the oxidative stress (low or high H_2_O_2_), we corrected for the general decrease in protein synthesis by normalizing the heavy fraction of each peptide in treated samples over the average fold-change between the treated and control samples of all peptides. Subsequently, we used the linear mixed effect model implemented in MSqRob (Goeminne et al., 2018) to calculate the statistical significance of the differential incorporation between H_2_O_2_-treated and control samples. The treatment was set as fixed effect of interest, the different peptides of the same protein as random effects, and protein name as grouping factor. Half of the interquantile range of the average difference between the normalized treated samples and the controls was used as minimal difference for a comparison to be accepted as significant. To correct for multiple testing, Benjamini-Hochberg correction was applied with an FDR cut-off < 0.01 (Table S4 and 5). Protein heavy fractions were determined by combining the peptide-specific heavy fractions by their median value and are reported in Table S3. To visualize the fold change relative to control (in Figure 5c), each protein heavy fraction was normalized over the median of the average observed values in the control samples.

To analyze the protein composition of ribosomes across different translational states, an available polysome proteome profiling dataset was downloaded ((Imami et al., 2018), PRIDE: PXD009292). The abundance of ribosomal proteins in selected fractions (40S, 60S, 80S and polysome) were extracted and the value of each ribosomal protein was normalized over the median abundance of all proteins of the corresponding subunit within each fraction (when considering proteins of the small subunit, the fraction corresponding to the 60S was excluded, and vice versa for the large subunit and the 40S fraction). For unsupervised clustering, the normalized levels of ribosomal proteins across fractions were hierarchically clustered using Euclidean distance, and clusters were visualized using the pheatmap R-package (RRID: SCR_016418, https://CRAN.R-project.org/package=pheatmap).

#### Structural analysis

The surface accessible areas of ribosomal proteins were calculated using the PDBePISA web service of the EBI (PDBe PISA v1.52 [20/10/2014], https://www.ebi.ac.uk/pdbe/pisa/pistart.html, Krissinel et Henrick, 2007) using the structure of small and large subunits of the human ribosome (PDB entry: 4V6X, https://www.rcsb.org/sequence/4V6X, (Anger et al., 2013)). In brief, PDB data were imported using the biological assembly CIF file, and total surface and interface areas were calculated for each chain using standard parameters in PISA. The solvent-accessible surface areas (calculated from the total surface area and all interface areas including RPs and rRNAs) values were extracted and summed up for every chain in the dataset.

#### Detection of newly synthesized rRNA in assembled ribosomes

New RNA was labeled by incubating cells for 3 hrs with 5 mM EU (5-Ethynyl Uridine) (ThermoFisher, E10345). Assembled ribosomes were purified by sucrose cushioning (see above) and the ribosome pellet was resuspended in 1 mL of TRIzol (ThermoFisher, 15596018). New RNA was purified by Click-iT Nascent RNA Capture Kit (ThermoFisher, C10365), according to the manufacturer’s recommendations. Briefly, 500 ng of RNA were clicked with 0.5 mM Biotin-Azide and EU-labeled RNA was purified by Dnyabeads MyOne Streaptavidin T1 beads. After washes, 1 µL of pre-diluted 1:200 ERCC RNA spike-in control mixes (ThermoFisher, 4456740) was added to all samples and reverse transcription was performed on the beads. The cDNA was then quantified by qPCR using TaqMan assay for the 18S rRNA (Thermo-fisher, Mm04277571_s1) and the ERCC-130 (5’-/5HEX/CGGAACAGG/ZEN/GCTGACGCCGC/3IANkFQ/-3’). To correct for differences in reverse transcription efficiency, each sample was internally normalized over ERCC-130 values. Finally, each experiment was normalized to a non-EU-labeled control.

#### Live cell imaging of RPL10a-PA-RFP

Cultured neurons were transfected at DIV 11-14, using the Magnetofectamine O2tm system, to express RPL10a tagged with a photoactivatable RFP (p323-L10A-PATagRFP, addgene plasmid #74172) and GFP as cell fill (pAcGFP1-C1, Clontech 632470). One or two days after transfection, neurons were imaged in supplemented E4 buffer (10 mM HEPES, 120 mM NaCl, 3 mM KCl, 10 mM D-Glucose, 2mM MgSO4 and 2 mM MgSO4, 1x B27, 1x GlutaMax, 1x MEM amino acids) with an inverted spinning disk confocal microscope (Zeiss 3i imaging systems; model CSU-X1). Images were acquired with Plan-Apochromat 63x/1.4 Oil DIC objective, at 488 nm (5 mW laser power and 50 ms exposure) and 561 nm (20 to 30 mW laser power and 100 ms exposure), using the Slidebook 5.5 software. Transfected cells were identified as GFP-positive. Z-stack (with 0.63 µm increments) was set to cover the whole cell body, and time-lapse was set with 30 min intervals. Photoactivation of a circular region around the nucleolus (identified by the lack of GFP signal) was performed using the 445 nm laser at 100 mW (two repetitions of 5 ms, at the center of the z plane). The sum intensity projection was used for image analysis. After background noise removal, the intensity inside the whole nucleus was quantified at each time point, and normalized over the maximum intensity reached after photoactivation.

#### Western blot

Samples (total cell lysates or sucrose cushion) were prepared as described above. After addition of NuPAGE LDS Sample Buffer (ThermoFisher, NP007) and NuPAGe Sample Reducing Agent (ThermoFisher, NP004) to a final concentration of 1x, samples were loaded onto 4% to 12% Bis-Tris NuPAGE gels (ThermoFisher). Gels were transferred using the Trans-Blot Turbo Transfer Pack (Biorad, 1704157) on a PVDF membrane (Immobilon-FL, IPFL00010 0.45 µm pore size). Membranes were stained using Revert 700 Total Protein Stain (LI-COR, 926-11015) for loading normalization. Immunoblotting was performed with primary antibodies as indicated (see table below for antibodies information) and secondary antibodies IRDye 680 and 800 (1:5000, LI-COR 926-68071, 926-68020, 926-32210, 926-32211). Images were acquired using LI-COR Image Studio Lite (RRID:SCR_013715) and analyzed using ImageJ/FIJI.

#### FISH Probe set

**FISH Probe set:**
| **Protein name** | **Gene symbol** | **ViewRNA ISH Cell Assay Probes set (ThermoFisher, QVC0001)** |
| --- | --- | --- |
| CamKIIa | CaMK2a | VC1-14332 |
| Histone3 | H3f3b | VC6-11342 |
| RACK1 | Gnbl2l1 | VC1-3061703 |
| RPL10/uL16 | Rpl10 | VC1-20759 |
| RPL13a/uL13 | Rpl13a | VC6-3229187 |
| RPL18a/eL20 | Rpl18a | VC6-3234577 |
| RPL19/eL19 | Rpl19 | VC1-17134 |
| RPL22/eL22 | Rpl22 | VC1-3061648 |
| RPL26/uL24 | Rpl26 | VC1-20756 |
| RPL29/eL29 | Rpl29 | VC1-20757 |
| RPL36/eL36 | Rpl36 | VC1-3060810 |
| RPL36A/eL42 | Rpl36a | VC1-3065218 |
| RPL38/eL38 | Rpl38 | VC1-3077564 |
| RPL4/uL4 | Rpl4 | VC1-20758 |
| RPL5/uL18 | Rpl5 | VC6-3231617 |
| RPL7a/eL8 | Rpl7a | VC6-3236240 |
| RPLP0/uL10 | Rplp0 | VC1-10192 |
| RPLP1/P1 | Rplp1 | VC1-3062658 |
| RPLP2/P2 | Rplp2 | VC1-3062659 |
| RPS11/uS17 | Rps11 | VC1-3061654 |
| RPS13/uS15 | Rps13 | VC1-20760 |
| RPS15a/uS8 | Rps15a | VC1-3062503 |
| RPS17/eS17 | Rps17 | VC1-3060264 |
| RPS21/eS21 | Rps21 | VC1-3061655 |
| RPS25/eS25 | Rps25 | VC1-3062623 |
| RPS26/eS26 | Rps26 | VC1-3060146 |
| RPS27/eS27 | Rps27 | VC1-3062104 |
| RPS29/uS14 | Rps29 | VC1-10529 |
| RPS30/eS30 | Fau | VC1-3060571 |
| RPS6/eS6 | Rps6 | VC1-3060274 |
| RPS7/eS7 | Rps7 | VC6-3060244 |

#### Antibody list

**Antibody list:**
| **Target** | **Company** | **Identifier** | **Dilution** |
| --- | --- | --- | --- |
| GAPDH | abcam | ab8245 | 1:2500 (WB) |
| MAP2 | SYSY | 188004 | 1:2000 or<br>1:1000 (IF) |
| Nucleolin | abcam | ab31163 | 1:10000 (IF)<br>1:2000 (WB) |
| puromycin | Kerafast | EQ0001 | 1:3500 (IF) |
| puromycin | CRB-cambridge | RANV10RbE76 | 1:200 (IF) |
| RACK1 | abcam | ab62735 | 1:200 (IF) |
| RanBP1 | abcam | ab97659 | 1:200 (IF) |
| RPL19/eL19 | abcam | ab224592 | 1:200 (IF) |
| RPL23/uL14 | Proteintech | 16086-1-AP | 1:200 (IF) |
| RPL26/uL24 | SIGMA | R0655 | 1:1000 (IF) |
| RPL36A/eL42 | Santa Cruz | sc-100831 | 1:200 (IF) |
| RPL38/eL38 | Bethyl | A305-412A | 1:2000 (IF) |
| RPL5/uL18 | abcam | ab186857 | 1:1000 (WB) |
| RPS11/uS17 | Bethyl | A303-936A | 1:1000 (IF)<br>1:1000 (WB) |
| RPS15/uS19 | abcam | ab154936 | 1:500 (IF) |
| RPS25/eS25 | ThermoFisher | PA5-56865 | 1:1000 (IF) |
| RPS26/eS26 | abcam | ab229571 | 1:50 (IF) |
| RPS28/eS28 | abcam | ab133963 | 1:50 (IF) |
| RPS3/uS3 | Bethyl | A303-840A-M | 1:1000 (WB)<br>1:100 (IF) |
| RPS30/eS30 | abcam | ab239073 | 1:500 (IF) |
| RPS3A/eS1 | Bethyl | A305-001A | 1:1000 (IF) |
| RPS5/uS7 | Bethyl | A304-010A-M | 1:250 (IF) |
| RPS9/uS4 | Bethyl | A303-946A-M | 1:400 (IF) |
| Y10b | abcam | ab171119 | 1:1000 (IF) |
| Goat anti-guinea pig<br>Dylight405 | Jackson<br>ImmunoResearch | 106-475-003 | 1:1000 (IF) |
| Goat anti-guinea pig-Alexa488 | ThermoFisher | A11073 | 1:1000 (IF) |
| Goat anti-mouse-Alexa594 | ThermoFisher | A11005 | 1:1000 (IF) |
| Goat anti-mouse-Alexa488 | ThermoFisher | A11001 | 1:1000 (IF) |
| Goat anti-rabbit-Alexa594 | ThermoFisher | A11037 | 1:1000 (IF) |
| Goat anti- rabbit-<br>Alexa488 | ThermoFisher | A11008 | 1:1000 (IF) |

## References

Aakalu, G., Smith, W.B., Nguyen, N., Jiang, C., Schuman, E.M., 2001. Dynamic visualization of local protein synthesis in hippocampal neurons. Neuron 30, 489–502. doi:10.1016/s0896-6273(01)00295-1

Alvarez-Castelao, B., tom Dieck, S., Fusco, C.M., Donlin-Asp, P., Perez, J.D., Schuman, E.M., 2020. The switch-like expression of heme-regulated kinase 1 mediates neuronal proteostasis following proteasome inhibition. eLife 9. doi:10.7554/eLife.52714

An, H., Harper, J.W., 2019. Ribosome Abundance Control Via the Ubiquitin-Proteasome System and Autophagy. Journal of Molecular Biology 432, 1–15. doi:10.1016/j.jmb.2019.06.001

Andreassi, C., Zimmermann, C., Mitter, R., Fusco, S., De Vita, S., Devita, S., Saiardi, A., Riccio, A., 2010. An NGF-responsive element targets myo-inositol monophosphatase-1 mRNA to sympathetic neuron axons. Nat Neurosci 13, 291–301. doi:10.1038/nn.2486

Anger, A.M., Armache, J.-P., Berninghausen, O., Habeck, M., Subklewe, M., Wilson, D.N., Beckmann, R., 2013. Structures of the human and Drosophila 80S ribosome. Nature 497, 80–85. doi:10.1038/nature12104

Biever, A., Glock, C., Tushev, G., Ciirdaeva, E., Dalmay, T., Langer, J.D., Schuman, E.M., 2020. Monosomes actively translate synaptic mRNAs in neuronal processes. Science 367, eaay4991–16. doi:10.1126/science.aay4991

Bodian, D., 1965. A suggestive relationship of nerve cell RNA with specific synaptic sites. Proc Natl Acad Sci USA 53, 418–425. doi:10.1073/pnas.53.2.418

Bogenhagen, D.F., Ostermeyer-Fay, A.G., Haley, J.D., Garcia-Diaz, M., 2018. Kinetics and Mechanism of Mammalian Mitochondrial Ribosome Assembly. CellReports 22, 1935–1944. doi:10.1016/j.celrep.2018.01.066

Briese, M., Saal, L., Appenzeller, S., Moradi, M., Baluapuri, A., Sendtner, M., 2016. Whole transcriptome profiling reveals the RNA content of motor axons. Nucleic Acids Res. 44, e33. doi:10.1093/nar/gkv1027

Burgin, K.E., Waxham, M.N., Rickling, S., Westgate, S.A., Mobley, W.C., Kelly, P.T., 1990. In situ hybridization histochemistry of Ca2+/calmodulin-dependent protein kinase in developing rat brain. J Neurosci 10, 1788–1798. doi:10.1523/JNEUROSCI.10-06-01788.1990

Cagnetta, R., Frese, C.K., Shigeoka, T., Krijgsveld, J., Holt, C.E., 2018. Rapid Cue-Specific Remodeling of the Nascent Axonal Proteome. Neuron 99, 29–46.e4. doi:10.1016/j.neuron.2018.06.004

Cajigas, I.J., Tushev, G., Will, T.J., tom Dieck, S., Fuerst, N., Schuman, E.M., 2012. The Local Transcriptome in the Synaptic Neuropil Revealed by Deep Sequencing and High-Resolution Imaging. Neuron 74, 453–466. doi:10.1016/j.neuron.2012.02.036

Cox, J., Mann, M., 2008. MaxQuant enables high peptide identification rates, individualized p.p.b.-range mass accuracies and proteome-wide protein quantification. Nature Biotechnology 26, 1367–1372. doi:10.1038/nbt.1511

Dörrbaum, A.R., Kochen, L., Langer, J.D., Schuman, E.M., 2018. Local and global influences on protein turnover in neurons and glia. eLife 7, 489. doi:10.7554/eLife.34202

Emmott, E., Jovanovic, M., Slavov, N., 2018. Ribosome Stoichiometry: From Form to Function. Trends Biochem Sci 44, 1–15. doi:10.1016/j.tibs.2018.10.009

Ferretti, M.B., Ghalei, H., Ward, E.A., Potts, E.L., Karbstein, K., 2017. Rps26 directs mRNA-specific translation by recognition of Kozak sequence elements 24, 1–13. doi:10.1038/nsmb.3442

Fornasiero, E.F., Mandad, S., Wildhagen, H., Alevra, M., Rammner, B., Keihani, S., Opazo, F., Urban, I., Ischebeck, T., Sakib, M.S., Fard, M.K., Kirli, K., Centeno, T.P., Vidal, R.O., Rahman, R.-U., Benito, E., Fischer, A., Dennerlein, S., Rehling, P., Feussner, I., Bonn, S., Simons, M., Urlaub, H., Rizzoli, S.O., 2018. Precisely measured protein lifetimes in the mouse brain reveal differences across tissues and subcellular fractions. Nat Comms 9, 4230. doi:10.1038/s41467-018-06519-0

Genuth, N.R., Barna, M., 2018. The Discovery of Ribosome Heterogeneity and Its Implications for Gene Regulation and Organismal Life. Mol Cell 71, 364–374. doi:10.1016/j.molcel.2018.07.018

Gioio, A.E., Lavina, Z.S., Jurkovicova, D., Zhang, H., Eyman, M., Giuditta, A., Kaplan, B.B., 2004. Nerve terminals of squid photoreceptor neurons contain a heterogeneous population of mRNAs and translate a transfected reporter mRNA. Eur J Neurosci 20, 865–872. doi:10.1111/j.1460-9568.2004.03538.x

Goeminne, L.J.E., Gevaert, K., Clement, L., 2018. Experimental design and data-analysis in label-free quantitative LC/MS proteomics: A tutorial with MSqRob. Journal of Proteomics 171, 23–36. doi:10.1016/j.jprot.2017.04.004

Granneman, S., Tollervey, D., 2007. Building Ribosomes: Even More Expensive Than Expected? Current Biology 17, R415–R417. doi:10.1016/j.cub.2007.04.011

Grimm, A., Cummins, N., Götz, J., 2018. Local Oxidative Damage in the Soma and Dendrites Quarantines Neuronal Mitochondria at the Site of Insult. iScience 6, 114–127. doi:10.1016/j.isci.2018.07.015

Gumy, L.F., Yeo, G.S.H., Tung, Y.-C.L., Zivraj, K.H., Willis, D., Coppola, G., Lam, B.Y.H., Twiss, J.L., Holt, C.E., Fawcett, J.W., 2011. Transcriptome analysis of embryonic and adult sensory axons reveals changes in mRNA repertoire localization. RNA 17, 85–98. doi:10.1261/rna.2386111

Hafner, A.-S., Donlin-Asp, P.G., Leitch, B., Herzog, E., Schuman, E.M., 2019. Local protein synthesis is a ubiquitous feature of neuronal pre- and postsynaptic compartments. Science 364, eaau3644. doi:10.1126/science.aau3644

Holt, C.E., Martin, K.C., Schuman, E.M., 2019. Local translation in neurons: visualization and function. 26, 557–566. doi:10.1038/s41594-019-0263-5

Imami, K., Milek, M., Bogdanow, B., Yasuda, T., Kastelic, N., Zauber, H., Ishihama, Y., Landthaler, M., Selbach, M., 2018. Phosphorylation of the Ribosomal Protein RPL12/ uL11 Affects Translation during Mitosis. Mol Cell 72, 84–98.e9. doi:10.1016/j.molcel.2018.08.019

Johnson, A.G., Lapointe, C.P., Wang, J., Corsepius, N.C., Choi, J., Fuchs, G., Puglisi, J.D., 2019. RACK1 on and off the ribosome. RNA 25, 881–895. doi:10.1261/rna.071217.119

Kang, K.R., Lee, S.-Y., 2001. Effect of serum and hydrogen peroxide on the Ca2+/calmodulin-dependent phosphorylation of eukaryotic elongation factor 2(eEF-2) in Chinese hamster ovary cells. Exp. Mol. Med. 33, 198–204. doi:10.1038/emm.2001.33

la Cruz, de, J., Karbstein, K., Woolford, J.L., Jr., 2015. Functions of Ribosomal Proteins in Assembly of Eukaryotic Ribosomes In Vivo. Annu. Rev. Biochem. 84, 93–129. doi:10.1146/annurev-biochem-060614-033917

Larburu, N., Montellese, C., O’Donohue, M.-F., Kutay, U., Gleizes, P.-E., Plisson-Chastang, C., 2016. Structure of a human pre-40S particle points to a role for RACK1 in the final steps of 18S rRNA processing. Nucleic Acids Res. 44, 8465–8478. doi:10.1093/nar/gkw714

MacLean, B., Tomazela, D.M., Shulman, N., Chambers, M., Finney, G.L., Frewen, B., Kern, R., Tabb, D.L., Liebler, D.C., MacCoss, M.J., 2010. Skyline: an open source document editor for creating and analyzing targeted proteomics experiments. Bioinformatics 26, 966–968. doi:10.1093/bioinformatics/btq054

Majzoub, K., Hafirassou, M.L., Meignin, C., Goto, A., Marzi, S., Fedorova, A., Verdier, Y., Vinh, J., Hoffmann, J.A., Martin, F., Baumert, T.F., Schuster, C., Imler, J.-L., 2014. RACK1 Controls IRES-Mediated Translation of Viruses. Cell 159, 1086–1095. doi:10.1016/j.cell.2014.10.041

Mardakheh, F.K., Paul, A., Kümper, S., Sadok, A., Paterson, H., Mccarthy, A., Yuan, Y., Marshall, C.J., 2015. Global Analysis of mRNA, Translation, and Protein Localization: Local Translation Is a Key Regulator of Cell Protrusions. Developmental Cell 35, 344–357. doi:10.1016/j.devcel.2015.10.005

Massaad, C.A., Klann, E., 2011. Reactive oxygen species in the regulation of synaptic plasticity and memory. Antioxidants & Redox Signaling 14, 2013–2054. doi:10.1089/ars.2010.3208

Mazaré, N., Oudart, M., Moulard, J., Cheung, G., Tortuyaux, R., Mailly, P., Mazaud, D., Bemelmans, A.-P., Boulay, A.-C., Blugeon, C., Jourdren, L., Le Crom, S., Rouach, N., Cohen-Salmon, M., 2020. Local Translation in Perisynaptic Astrocytic Processes Is Specific and Changes after Fear Conditioning 32, 108076–22. doi:10.1016/j.celrep.2020.108076

McGlincy, N.J., Ingolia, N.T., 2017. Transcriptome-wide measurement of translation by ribosome profiling. Methods 126, 112–129. doi:10.1016/j.ymeth.2017.05.028

Middleton, S.A., Eberwine, J., Kim, J., 2019. Comprehensive catalog of dendritically localized mRNA isoforms from sub-cellular sequencing of single mouse neurons. BMC Biol. 17, 5. doi:10.1186/s12915-019-0630-z

Miller, S., Yasuda, M., Coats, J.K., Jones, Y., Martone, M.E., Mayford, M., 2002. Disruption of dendritic translation of CaMKIIalpha impairs stabilization of synaptic plasticity and memory consolidation. Neuron 36, 507–519. doi:10.1016/s0896-6273(02)00978-9

Mirzaei, H., Regnier, F., 2007. Identification of yeast oxidized proteins. Journal of Chromatography A 1141, 22–31. doi:10.1016/j.chroma.2006.11.009

Mishchenko, Y., Hu, T., Spacek, J., Mendenhall, J., Harris, K.M., Chklovskii, D.B., 2010. Ultrastructural analysis of hippocampal neuropil from the connectomics perspective. Neuron 67, 1009–1020. doi:10.1016/j.neuron.2010.08.014

Misra, M., Edmund, H., Ennis, D., Schlueter, M.A., Marot, J.E., Tambasco, J., Barlow, I., Sigurbjornsdottir, S., Mathew, R., Vallés, A.M., Wojciech, W., Roth, S., Davis, I., Leptin, M., Gavis, E.R., 2016. A Genome-Wide Screen for Dendritically Localized RNAs Identifies Genes Required for Dendrite Morphogenesis. G3 6, 2397–2405. doi:10.1534/g3.116.030353

Moccia, R., Chen, D., Lyles, V., Kapuya, E., E, Y., Kalachikov, S., Spahn, C.M.T., Frank, J., Kandel, E.R., Barad, M., Martin, K.C., 2003a. An unbiased cDNA library prepared from isolated Aplysia sensory neuron processes is enriched for cytoskeletal and translational mRNAs. Journal of Neuroscience 23, 9409–9417. doi:10.1523/JNEUROSCI.23-28-09409.2003

Moccia, R., Chen, D., Lyles, V., Kapuya, E., E, Y., Kalachikov, S., Spahn, C.M.T., Frank, J., Kandel, E.R., Barad, M., Martin, K.C., 2003b. An unbiased cDNA library prepared from isolated Aplysia sensory neuron processes is enriched for cytoskeletal and translational mRNAs. Journal of Neuroscience 23, 9409–9417. doi:10.1523/JNEUROSCI.23-28-09409.2003

Moor, A.E., Golan, M., Massasa, E.E., Lemze, D., Weizman, T., Shenhav, R., Baydatch, S., Mizrahi, O., Winkler, R., Golani, O., Stern-Ginossar, N., Itzkovitz, S., 2017. Global mRNA polarization regulates translation efficiency in the intestinal epithelium. Science 357, 1299–1303. doi:10.1126/science.aan2399

Perez, J.D., Dieck, S.T., Alvarez-Castelao, B., Tushev, G., Chan, I.C., Schuman, E.M., 2021. Subcellular sequencing of single neurons reveals the dendritic transcriptome of GABAergic interneurons. eLife 10. doi:10.7554/eLife.63092

Peterson, A.C., Russell, J.D., Bailey, D.J., Westphall, M.S., Coon, J.J., 2012. Parallel reaction monitoring for high resolution and high mass accuracy quantitative, targeted proteomics. Mol Cell Proteomics 11, 1475–1488. doi:10.1074/mcp.O112.020131

Pérez-Riverol, Y., Csordas, A., Bai, J., Bernal-Llinares, M., Hewapathirana, S., Kundu, D.J., Inuganti, A., Griss, J., Mayer, G., Eisenacher, M., Pérez, E., Uszkoreit, J., Pfeuffer, J., Sachsenberg, T., Yilmaz, Ş., Tiwary, S., Cox, J., Audain, E., Walzer, M., Jarnuczak, A.F., Ternent, T., Brazma, A., Vizcaíno, J.A., 2019. The PRIDE database and related tools and resources in 2019: Improving support for quantification data. Nucleic Acids Res. 47, D442–D450. doi:10.1093/nar/gky1106

Poon, M.M., Choi, S.-H., Jamieson, C.A.M., Geschwind, D.H., Martin, K.C., 2006. Identification of process-localized mRNAs from cultured rodent hippocampal neurons. Journal of Neuroscience 26, 13390–13399. doi:10.1523/JNEUROSCI.3432-06.2006

Poulopoulos, A., Murphy, A.J., Ozkan, A., Davis, P., Hatch, J., Kirchner, R., Macklis, J.D., 2019. Subcellular transcriptomes and proteomes of developing axon projections in the cerebral cortex. Nature 565, 356–360. doi:10.1038/s41586-018-0847-y

Pulk, A., Liiv, A., Peil, L., Maiväli, Ü., Nierhaus, K., Remme, J., 2010. Ribosome reactivation by replacement of damaged proteins. Mol. Microbiol. 75, 801–814. doi:10.1111/j.1365-2958.2009.07002.x

Rappsilber, J., Mann, M., Ishihama, Y., 2007. Protocol for micro-purification, enrichment, pre-fractionation and storage of peptides for proteomics using StageTips. Nature protocols 2, 1896–1906. doi:10.1038/nprot.2007.261

Ross, A.B., Langer, J.D., Jovanovic, M., 2021. Proteome Turnover in the Spotlight: Approaches, Applications, and Perspectives. Mol Cell Proteomics 20, 100016. doi:10.1074/mcp.R120.002190

Saal, L., Briese, M., Kneitz, S., Glinka, M., Sendtner, M., 2014. Subcellular transcriptome alterations in a cell culture model of spinal muscular atrophy point to widespread defects in axonal growth and presynaptic differentiation. RNA 20, 1789–1802. doi:10.1261/rna.047373.114

Schwanhäusser, B., Gossen, M., Dittmar, G., Selbach, M., 2009. Global analysis of cellular protein translation by pulsed SILAC. Proteomics 9, 205–209. doi:10.1002/pmic.200800275

Shenton, D., Smirnova, J.B., Selley, J.N., Carroll, K., Hubbard, S.J., Pavitt, G.D., Ashe, M.P., Grant, C.M., 2006. Global Translational Responses to Oxidative Stress Impact upon Multiple Levels of Protein Synthesis*. J Biol Chem 281, 29011–29021. doi:10.1074/jbc.M601545200

Shi, Z., Fujii, K., Kovary, K.M., Genuth, N.R., Röst, H.L., Teruel, M.N., Barna, M., 2017. Heterogeneous Ribosomes Preferentially Translate Distinct Subpools of mRNAs Genome-wide. Mol Cell 67, 71–83.e7. doi:10.1016/j.molcel.2017.05.021

Shigeoka, T., Jung, H., Jung, J., Turner-Bridger, B., Ohk, J., Lin, J.Q., Amieux, P.S., Holt, C.E., 2016. Dynamic Axonal Translation in Developing and Mature Visual Circuits. Cell 166, 181–192. doi:10.1016/j.cell.2016.05.029

Shigeoka, T., Koppers, M., Wong, H.H.-W., Lin, J.Q., Cagnetta, R., Dwivedy, A., de Freitas Nascimento, J., van Tartwijk, F.W., Ströhl, F., Cioni, J.-M., Schaeffer, J., Carrington, M., Kaminski, C.F., Jung, H., Harris, W.A., Holt, C.E., 2019. On-Site Ribosome Remodeling by Locally Synthesized Ribosomal Proteins in Axons. CellReports 29, 3605–3619.e10. doi:10.1016/j.celrep.2019.11.025

Slavov, N., Semrau, S., Airoldi, E., Budnik, B., van Oudenaarden, A., 2015. Differential Stoichiometry among Core Ribosomal Proteins 13, 865–873. doi:10.1016/j.celrep.2015.09.056

Stoykova, A.S., Dudov, K.P., Dabeva, M.D., Hadjiolov, A.A., 1983. Different rates of synthesis and turnover of ribosomal RNA in rat brain and liver. J Neurochem 41, 942–949. doi:10.1111/j.1471-4159.1983.tb09038.x

Taylor, A.M., Berchtold, N.C., Perreau, V.M., Tu, C.H., Li Jeon, N., Cotman, C.W., 2009. Axonal mRNA in uninjured and regenerating cortical mammalian axons. Journal of Neuroscience 29, 4697–4707. doi:10.1523/JNEUROSCI.6130-08.2009

Tennyson, V.M., 1970. The fine structure of the axon and growth cone of the dorsal root neuroblast of the rabbit embryo. Journal of Cell Biology 44, 62–79. doi:10.1083/jcb.44.1.62

Thomas, F., Kutay, U., 2003. Biogenesis and nuclear export of ribosomal subunits in higher eukaryotes depend on the CRM1 export pathway. Journal of Cell Science 116, 2409–2419. doi:10.1242/jcs.00464

tom Dieck, S., Kochen, L., Hanus, C., Heumüller, M., Bartnik, I., Nassim-Assir, B., Merk, K., Mosler, T., Garg, S., Bunse, S., Tirrell, D.A., Schuman, E.M., 2015. Direct visualization of newly synthesized target proteins in situ. Nature Methods 12, 411–414. doi:10.1038/nmeth.3319

Tsurugi, K., Ogata, K., 1985. Evidence for the exchangeability of acidic ribosomal proteins on cytoplasmic ribosomes in regenerating rat liver. Journal of Biochemistry 98, 1427–1431. doi:10.1093/oxfordjournals.jbchem.a135410

Tushev, G., Glock, C., Heumüller, M., Biever, A., Jovanovic, M., Schuman, E.M., 2018. Alternative 3’ UTRs Modify the Localization, Regulatory Potential, Stability, and Plasticity of mRNAs in Neuronal Compartments. Neuron 98, 495–511.e6. doi:10.1016/j.neuron.2018.03.030

Tyanova, S., Temu, T., Cox, J., 2016. The MaxQuant computational platform for mass spectrometry-based shotgun proteomics. Nature protocols 11, 2301–2319. doi:10.1038/nprot.2016.136

Warner, J.R., 1999. The economics of ribosome biosynthesis in yeast. Trends Biochem Sci 24, 437–440. doi:10.1016/s0968-0004(99)01460-7

Warner, J.R., Udem, S.A., 1972. Temperature sensitive mutations affecting ribosome synthesis in Saccharomyces cerevisiae. Journal of Molecular Biology 65, 243–257. doi:10.1016/0022-2836(72)90280-x

Wiśniewski, J.R., Zougman, A., Nagaraj, N., Mann, M., 2009. Universal sample preparation method for proteome analysis. Nature Methods 6, 359–362. doi:10.1038/nmeth.1322

Wu, C.C.-C., Zinshteyn, B., Wehner, K.A., Green, R., 2019. High-Resolution Ribosome Profiling Defines Discrete Ribosome Elongation States and Translational Regulation during Cellular Stress. Mol Cell 73, 959–970.e5. doi:10.1016/j.molcel.2018.12.009

Xue, S., Tian, S., Fujii, K., Kladwang, W., Das, R., Barna, M., 2015. RNA regulons in Hox 5’ UTRs confer ribosome specificity to gene regulation. Nature 517, 33–38. doi:10.1038/nature14010

Zinker, A.B.-S.A.S., 2014. The P1/P2 Protein Heterodimers Assemble to the Ribosomal Stalk at the Moment When the Ribosome Is Committed to Translation but Not to the Native 60S Ribosomal Subunit in Saccharomyces cerevisiae. Biochemistry 53, 1–8. doi:10.1021/bi500341w

Zivraj, K.H., Tung, Y.-C.L., Piper, M., Gumy, L., Fawcett, J.W., Yeo, G.S.H., Holt, C.E., 2010. Subcellular profiling reveals distinct and developmentally regulated repertoire of growth cone mRNAs. Journal of Neuroscience 30, 15464–15478. doi:10.1523/JNEUROSCI.1800-10.2010

